# Emergence of active contractile patterns alters monolayer force generation and transmission in response to focal adhesion distribution

**DOI:** 10.1101/2024.04.10.588783

**Authors:** John Robert Davis, Josephine Solowiej-Wedderburn, Sebastián L. Vega, Jason A. Burdick, Carina Dunlop, Nicolas Tapon

## Abstract

For tissue development, cells must generate contractile forces which are transmitted to their surrounding matrix or neighbouring cells via adhesion complexes. It is often envisaged that a simple linear counterbalance of cell generated stress with extracellular matrix (ECM) traction forces exists. However, experimental evidence indicates that modulating cell-ECM attachment does not necessarily lead to expected reciprocal changes in intercellular stresses. As ECM composition or mechanical properties are rarely uniform, it is important to understand the complexity of how focal adhesions alter stress transmission and the force-balance of a tissue. To address this, we confined monolayers on adhesive patterns altering focal adhesion distribution. Traction force microscopy and laser ablations of cell-cell junctions were used to examine stresses across epithelial monolayers whilst modulating substrate stiffness. We show that monolayers reach different force-balance states depending on focal adhesion distribution. Using an active matter model and confirmed experimentally, we reveal that a force-balance is generated by non-uniform patterns of cell contractility linked to adhesion patterning. This work highlights the importance of integrating the position of cell-ECM attachments into our vision of the mechanical landscape of living tissues.

**Teaser:** To infer a tissue’s force-balance, positional information of focal adhesion distribution needs to be integrated due to the emergence of non-uniform patterns of cell contractility.

## Introduction

A key aspect of many developmental, physiological, and pathological processes is the ability of cells to generate and transmit forces across a tissue. Cells transmit intracellular stresses to their environment, and this is essential for orchestrating cooperative behaviours such as morphogenesis^1^, collective migration^2^ and wound healing^3^. Stresses are transmitted through adhesion complexes^4^; either to the substrate generating traction stresses, or to neighbouring cells generating intercellular stresses. Furthermore, these adhesion complexes are important mechanotransduction nodes regulating cellular processes such as metabolism^5^, gene transcription^6^, as well as actin architecture^7^. Therefore, to gain biological insight linking tissue mechanics to molecular dynamics, there is a need to quantify local traction stresses and intercellular stresses across epithelial monolayers.

Various methodologies are available to interrogate stresses within tissues^8^. Traction force microscopy, where the movement of microbeads embedded within a substrate are tracked, is a proven method for examining cell-generated forces transmitted to the substrate^9^. As a fundamental principle of tissue mechanics is the necessity for monolayers to balance traction and intercellular stresses to maintain epithelial integrity traction force microscopy has been extended to infer intercellular stresses based on mechanical arguments ^10–14^. This has been developed for a range of scenarios from cell doublets^14–17^, to small cell clusters^13^ to entire epithelial sheets^10,12,18,19^. A core assumption of these techniques is that intercellular stresses can be inferred from traction stresses through a passive elastic model, often diagrammatically evoked as a tug-of-war stalemate. This type of model leads to two conclusions: 1) that there is a linear relationship between the scalar sum of traction stresses and intercellular stresses within a monolayer^13,14^, and 2) that intercellular stress would be greatest in the centre of the monolayer and lowest at the boundary^20,21^. There are some experimental data based on Förster resonance energy transfer (FRET) tension sensors which support these conclusions. Both E-Cadherin and ZO-2 tension sensors have been reported to decrease in FRET efficiency (increase in molecular tension) in monolayers cultured on stiffer substrates^22,23^. Furthermore, E-Cadherin-α-catenin tension sensors have also been reported to exhibit a reduction in FRET efficiency away from the edge of a wounded epithelium^24^. However, it has also been reported in cell doublets that cell-cell forces inferred from traction force data do not correlate with the FRET-efficiency of an E-Cadherin tension sensor, with the authors proposing a molecular tension homeostasis mechanism to resolve these findings^16^. Furthermore, experiments examining intercellular stresses through laser ablation have also made observations which are not consistent with the conclusions drawn from passive elastic models linking traction stresses and intercellular stresses.

Laser ablation is a widely employed technique to quantify intercellular stresses within tissues and through rheological models infer tissue mechanical properties. This is achieved by examining the relaxation of cells after individual or groups of junctions have been cut^25,26^. By disrupting entire cell-cell junctions, this method provides a holistic overview of cell mechanics incorporating both cortical cytoskeleton and all cell-cell adhesion complexes. It is widely used in developmental systems that are not amenable to other types of biomechanical techniques to measure or infer tissue mechanics but is also applied to cells *in vitro* to examine both intracellular and intercellular stresses. Work in the *Drosophila* abdominal epidermis showed that degradation of the basal ECM occurring at the pupal stage led to an increase in intercellular tension^27^. Similarly, Madin-Darby canine kidney (MDCK) cells cultured on large micropatterned disks on soft hydrogels showed lower traction stresses but higher recoil velocities than those cultured on stiff hydrogels ^28^. These contradict the conclusions drawn from passive elastic models employed to link traction stresses to intercellular stresses. In both studies, intercellular tension was greatest when cell-ECM friction was reduced, in the case of *Drosophila* abdomen the ECM was degraded ^27^ and for MDCK cells there was weaker cell-ECM attachments on softer hydrogels^28^ which leads to a reduction in cell-ECM friction^29^. This supports pioneering work in amnioserosa cells during *Drosophila* embryonic dorsal closure showing that focal adhesions act as tethers that reduce both intercellular tension and stress transmission^30^. Taken together, these studies question the use of simple passive elastic models to infer intercellular stresses from traction stress which do not explicitly model the active cell contractile behaviours, leading to proposals to replace these types of models with active-matter models^31^. This approach directly accounts for active contractility within the force balance enabling both active cell-generated stress and the material stress generated in passive cell structures to be decoupled, and this approach is now widely adopted in a range of contexts^32–37^. The benefit of the active matter framework is that it is highly adaptable and capable of capturing a range of the complexity in the cell-ECM force balance including ECM stiffness-induced changes. Significantly, cell adhesions and adhesion patterning can be explicitly described^38,39^, which is significant given recent experimental findings that highlight that 1) the distribution of focal adhesions is important in determining the length-scale that stresses can be transmitted through the monolayer^30^, and 2) that the compliance of the substrate also influences the distance stresses can be transmitted through the substrate^40^. Indeed, it is intuitively clear that examining the relationship between intercellular and traction stresses across a monolayer requires an understanding of how both focal adhesion distribution and substrate compliance alter the acquired force-balance. This is especially true when considering that *in vivo* ECM composition, architecture and mechanical properties are often heterogeneous^41^.

To examine the relationship between traction and intercellular stresses we combined traction force microscopy and annular laser ablation^25^. As traction forces are known to be dependent on the size and geometry^34,38^ of cells and tissues, we employed micropatterning of hydrogels to standardise these factors. Furthermore, through micropatterning we modulated cell-ECM interactions by culturing monolayers on disks or rings to examine the role of focal adhesion distribution in determining monolayers mechanical equilibrium. We also modulated substrate compliance which is known to alter both substrate stress transmission^40^ and traction stresses generated by cells^38^. When focal adhesions are distributed throughout the monolayer, traction forces are dependent on the substrate stiffness within our experimentally tested range. However, when focal adhesion distribution is mainly restricted to the periphery, traction forces become independent of the substrate. Furthermore, we observe that intercellular stress is inversely proportional to substrate stiffness, so that intercellular stress reduces as hydrogel stiffness increases. We combined this experimental approach with an active stress model based on a continuum description, which explicitly modelled cell attachments and variation of cell activity within the layer. This revealed that the relationship between substrate stiffness and intercellular stress is elicited by the emergence of non-uniform contractility across the monolayer, with greater contractility present at regions with greater adhesions.

## Results

### Adhesive surface area dictates focal adhesion distribution across a monolayer

To examine the relationship between traction and intercellular stresses, we cultured non-tumorigenic mouse mammary epithelial cells (EpH4)^42^ on methacrylated hyaluronic acid (MeHA) hydrogels^43^. 3% w/v MeHA solutions were crosslinked with a range of dithiothreitol (DTT) crosslinker concentrations to modulate the rigidity of hydrogels from 10.8 ± 2.5 kPa to 31.4 ± 2.8 kPa at intervals of ∼5kPa (Fig. S1a-c). Whilst these values are higher than what has been reported for mouse mammary tissue^44^, they span reported basement membrane values^41^ and ranges employed in other systems examining stress transmission^14,28,29,40^. To examine the role of focal adhesions in determining monolayers mechanical equilibria, we modulated the adhesive surface area beneath monolayers by micropatterning laminin, EpH4 cell’s preferred ECM component^45^. MeHA hydrogels were patterned with either disk (Fig. 1a) or ring (Fig. 1b) micropatterns, with an outer diameter of 200μm (approximately 10 – 20 cells wide) and the width of rings being 20μm (approximately 1-2 cells wide). We conjugated an Alexa633 fluorophore to laminin to assess if increasing levels of MeHA crosslinking increased the concentration of laminin on the surface of hydrogels for both disk (Fig. 1c) and ring (Fig. 1d) patterns. Whilst there were fluctuations in laminin intensity on different hydrogel stiffnesses (Fig. 1c,d) these did not follow a particular pattern linked to substrate stiffness. Furthermore, whilst there were radial differences across the disk pattern (Fig. 1c), these did not correlate with differences in focal adhesions (Fig. 1g). Therefore, the concentration of laminin in patterned areas appears to not alter the adhesions that cells can generate with the substrate.

**Fig. 1.**
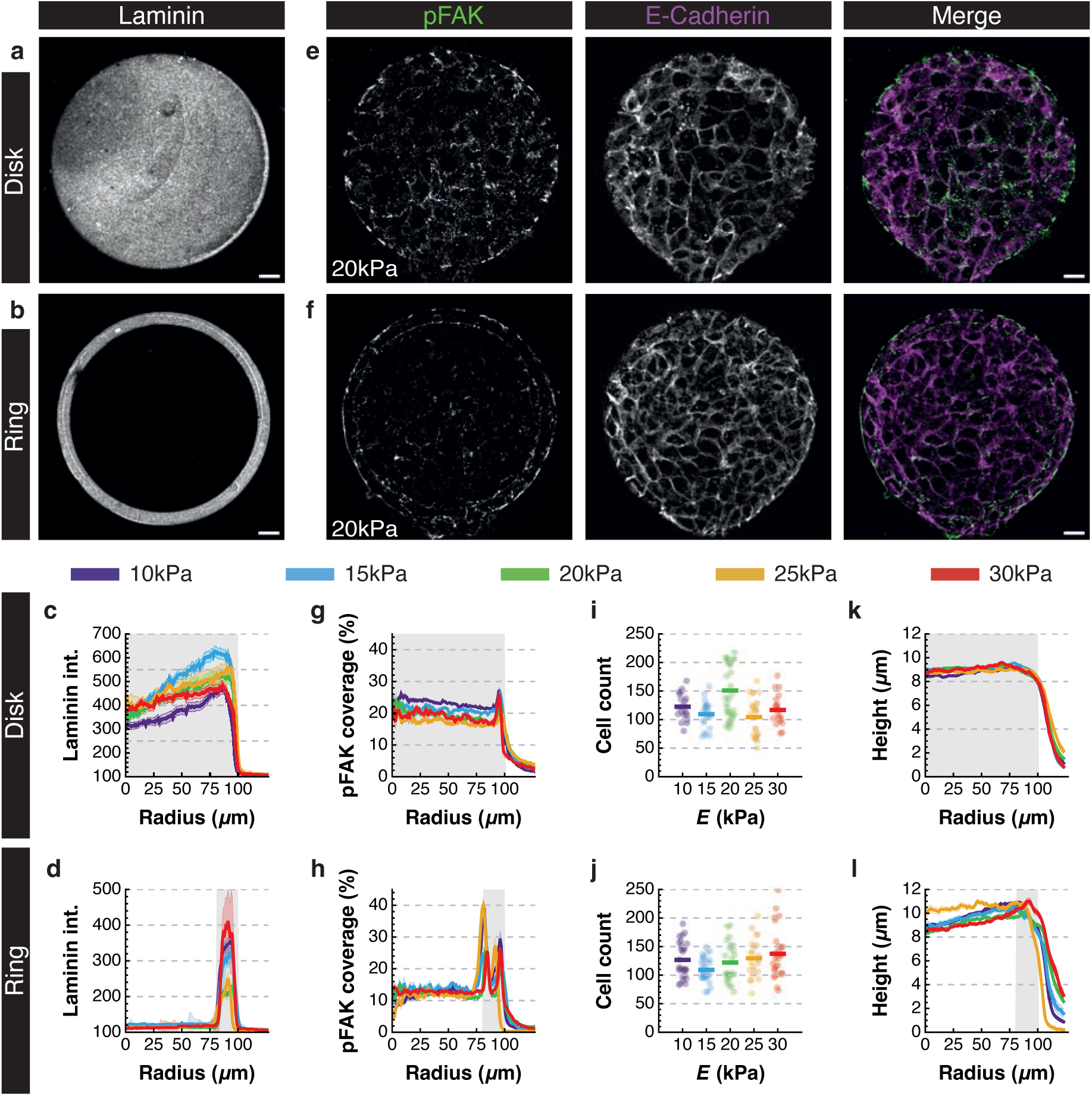
Modulating adhesive surface of EpH4 monolayers. **a,b** Example projection of laminin micropatterns for disk (**a**) and ring (**b**). **c,d** Quantification of mean laminin intensity radially across micropatterns. **e,f** Example projections of basal pFAK and apical E-Cadherin on disk (**e**) and ring (**f**) micropatterns. **g,h** Quantification of mean pFAK coverage radially across micropatterns. **i,j** Dot plot of cell number within each monolayer, bars highlight mean value for each population. **k,l** Quantification of mean cell height radially across micropatterns. For all quantification thick line are mean values and ribbons are S.E.M, grey shaded region represents adhesive surface. All N numbers and statistics included in Table S1 in Supplementary Information. All scale bars = 20μm.

To measure active focal adhesions on different patterns, we immunostained monolayers for phosphorylated focal adhesion kinase (pFAK) (Fig. 1e,f) as previously reported [add Pere paper]. Altering the adhesive surface did lead to changes in the distribution of pFAK across the monolayer. On disks, pFAK was evenly distributed across the entire monolayer with a slight peak at the periphery (Fig. 1g). On rings however, pFAK was mainly focused on the perimeter of the adhesive area with peaks at the boundaries, and a low-level present in the non-patterned area (Fig. 1h). As no laminin is present in the middle of the ring micropatterns (Fig. 1d), the low-level of pFAK staining in this region is probably due to nascent proteins secreted by EpH4 cells becoming bound to the substrate^46,47^ and not direct binding to the MeHA gel. Indeed, our hyaluronic modification rate was between 76 – 79% (Fig. S1a), far higher than the threshold for cells to bind to unmodified disaccharides^48^. Furthermore, whilst cells cultured on disks were surrounded by pFAK puncta (Fig. 1e) across the monolayer, on rings, cells in the centre tended to only have isolated puncta of pFAK (Fig. 1f). Examining the total pFAK coverage beneath disks and rings revealed that rings consistently had less total pFAK coverage than disks (Fig. S1d,e), suggesting that monolayers cultured on ring micropatterns had reduced active cell-ECM attachments. Surprisingly, given previous reports^29^ both patterns showed a slight reduction in total pFAK as substrate stiffness increased (Fig. S1d,e). The regulation of focal adhesion dynamics is dependent on many factors^49^, and there are contradictory reports on how substrate rigidity alters focal adhesion coverage^50,51^. It is not clear why total pFAK coverage reduces on stiffer hydrogels, however, cells have been reported to sense and respond to strain energy^52^, and as we are examining active focal adhesions it is possible that strain energy is decreasing as stiffness increases in EpH4 cells across the range of rigidities employed. In summary, we established experimental conditions that allow us to vary cell-ECM adhesion distribution and substrate stiffness.

We next examined if altering the adhesive pattern or substrate rigidity altered cell-cell adhesions or cell morphology across the monolayer. We observed no obvious differences in the localisation of E-Cadherin for both patterns on all hydrogel stiffnesses, with strong localisation along entire cell boundaries (Fig. 1e,f). Likewise, whilst there was a distribution in the number of cells that composed each monolayer, the distribution was similar for all patterns and substrate rigidities (Fig. 1i,j). Examining cell morphology, we observed that cell height does not significantly change as substrate rigidity increases (Fig. 1k,l). However, we did observe that whilst on disks height is relatively constant (8.9 ± 1.3μm; Fig. 1k), on rings height increases from 8.9 ± 1.8μm in the centre to 10.4 ± 1.8μm on the adhesive area (Fig. 1l). Cell area tended to be slightly greater in the centre of the patterns, especially for disks (Fig. S1f,g) and there tended to be a slight increase in elongation at the perimeter for both patterns (Fig. S1h,i). However, these variations were subtle between the different patterns suggesting that overall monolayer packing was similar between patterns and on all substrate stiffnesses.

Finally, we examined whether there were junctional rearrangements over the timescale of our movies. In our timelapse experiments, we imaged for one minute before performing laser ablations, so subtracted the frame prior to laser ablation from the initial frame and calculated the percentage pixel intensity of our membrane label that remained. This analysis revealed a value close to zero on both patterns and on all hydrogel stiffnesses (Fig. S1j,k). Furthermore, overlayed images of cell membranes for the same duration as our ablation experiments (5 minutes apart), showed a high degree of overlap (Fig. S1l). We therefore assume that junctional rearrangement is not a major contributing factor to the force balance dynamics of the monolayer at the timescale of our experiments.

### Focal adhesion patterning alters dependence of traction forces on substrate stiffness

We next wanted to assess how altering the adhesive area affected traction forces. To measure traction forces, we embedded fluorescent microbeads within the MeHA hydrogels and measured their displacement^53^. Following recent reporting of extensile forces generated by MDCK cell layers^28^, we first confirmed that all displacement vectors indicated contraction towards the centre of the cell layer across all substrates (Fig. 2a,b, Fig. S2a-d).

**Fig. 2.**
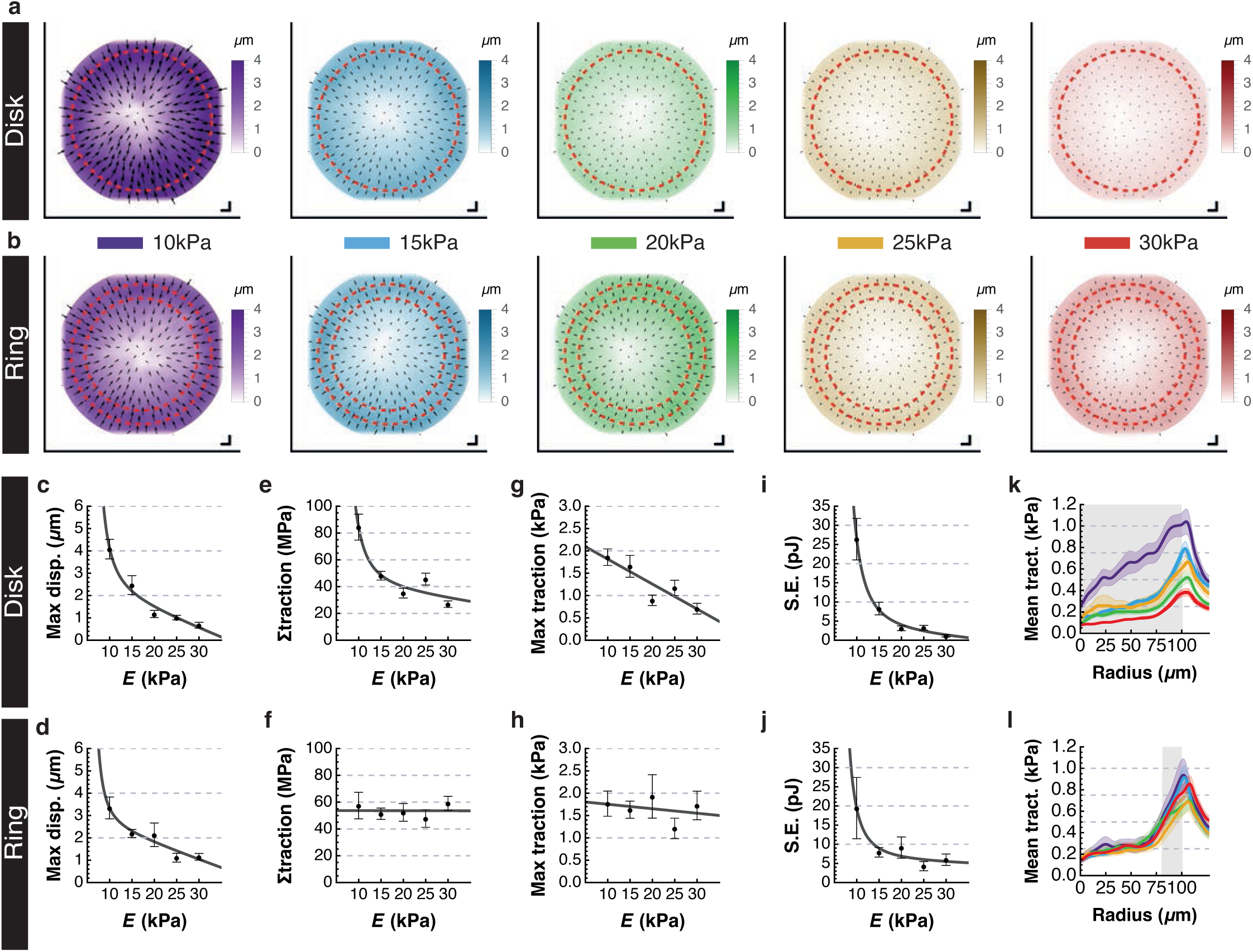
Monolayer traction stresses are altered by modulating adhesive surface. **a,b** Mean displacement vector plots for monolayers on disks (**a**) and rings (**b**) on different substrate stiffnesses. Red dashed line outlines the adhesive surface and scale bars represent 4μm. **c,d** Quantification of maximum bead displacement by monolayers adhered to disks and rings on different substrate stiffnesses. **e,f** Quantification of total traction forces generated by monolayers adhered to disks and rings on different substrate stiffnesses. **g,h** Quantification of maximum traction forces generated by monolayers adhered to disks and rings on different substrate stiffnesses. **i,j** Quantification of strain energy transferred by monolayers into substrate adhered to disks and rings on different substrate stiffnesses. For **c-j** black dots highlight mean value, whiskers represent S.E.M and grey line is line of best fit. **k,l** Quantification of mean traction forces radially across micropatterns. Thick lines are mean values and ribbons are S.E.M, grey shaded region represents adhesive surface.

We also observed the greatest displacements at the periphery of the monolayer (Fig. 2a,b, Fig. S2e,f). As expected from previous reports^28,38^, we observed a reduction in displacement magnitude as substrate rigidity increased for both disks and rings (Fig. 2c,d, Fig. S2g-h). We next examined the traction forces generated by the monolayers and noticed stark differences between disk and ring patterns (Fig. 2e-h). On disk patterns we observe a decrease in both the scalar sum of traction forces, which we term total traction, and the maximum traction force as substrate stiffness increased (Fig. 2e,g). On rings, total traction force remained almost constant (Fig. 2f), and maximum traction forces showed a weak reduction (Fig. 2h). It has been reported in micropatterned single MDCK cells that strain energy is not affected by substrate rigidity^38^. In the case of the monolayers considered here, as displacement decreased for both patterns and traction forces decreased on disks and remained constant on rings, we were interested in examining if our adhesive patterns showed marked differences in strain energy. We observed that for both disks and rings there is a decrease in strain energy as substrate stiffness increases, potentially explaining the reduction in pFAK. However, this difference is stronger for disks than it is for rings (Fig. 2i,j), with a mean strain energy reduction between 10kPa to 30kPa for disks (25,242pJ) being almost double that of rings (13,481pJ).

As the traction forces highlighted a clear difference in behaviour between disk and rings on different substrates, we decided to examine whether this reflected a difference in the spatial distribution of traction forces. Examining mean traction forces radially along the epithelial clusters showed the highest traction forces at the periphery for both ring and disk patterns on all substrates (Fig. 2k,l). Consistent with changes in total and maximum traction forces, for disks we observe a reduction in traction magnitude with increasing stiffness whilst on rings we observe virtually no change in traction force magnitude (Fig. 2k,l).

However, we noticed a slight change in the distribution of traction forces radially between the patterns, with traction forces appearing to be more localised at the periphery on rings than in disks especially on soft substrates (Fig. 2k,l). As our pFAK data shows on disks all cells are surrounded by focal adhesions, this suggests that on compliant substrate, cells within the monolayer locally contribute towards deforming the substrate, leading to a gradual radial accumulation in traction forces. However, as the hydrogel becomes more rigid and cells are less able to deform the substrate, traction forces become more localised to the edge, as previously theoretically predicted^54^. This is altered on ring patterns where the cells in the central non-adhesive region do not deform the substrate locally, so that the applied traction forces are always localised to the periphery. Taken together our results suggest that altering the adhesive surface leads to changes in the generation and distribution of traction forces across the epithelium.

### Intercellular stresses decrease with increasing substrate stiffness

To interrogate stress transmission within the monolayer, we combined theoretical and experimental approaches. The theoretical model specifically describes the monolayer as an active soft material in which the monolayer stresses and displacements are determined from continuum elasticity theory; specifically, we include an isotropic active stress in the relationship between stress and strain that accounts for the cell-derived contractility (Fig. 3a, see Supplemental Information Table 1 for model parameters) in contrast to passive models for traction-force inference. Such active matter models are widely adopted to describe active, contractile cells and tissues adhered to substrates^17,35,38,54^. Within our framework, substrate adhesion is coupled into the active model in the force balance equations through applied tractions, which may be restricted (as here) to act only where the layer is adhered^54^. Based on our experimental observations with pFAK staining (Fig. 1i,j), layers were either modelled as adhered uniformly to recapitulate the disk, or for the ring pattern the model assumes adhesions concentrated at the ring but with an accompanying underlying distribution of small adhesions sites everywhere else as observed (Fig. 3a). A more detailed description of the model can be found in supplemental information and methods section.

**Fig. 3.**
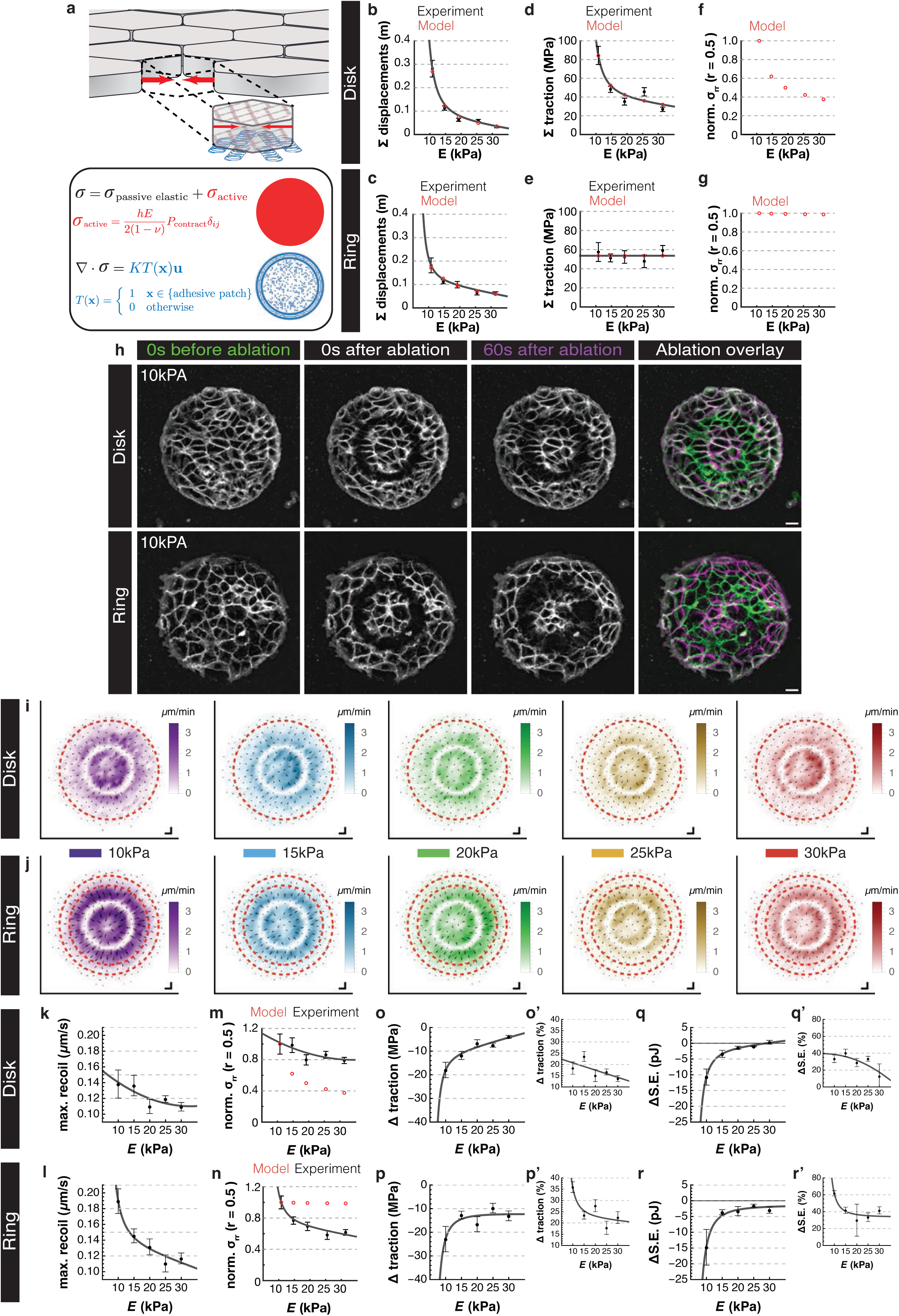
Intercellular stress is not proportional to traction stress. **a** Schematic representation of the major components that comprise the force balance model. Cells are considered as active contractile agents (red arrows) bound together to form a monolayer continuum as well as being bound to the substrate through adhesions (blue springs). Any stresses within the monolayer are therefore a product of these active forces and passive elastic forces transmitted through the monolayer (top equation). The active forces generated by the cells are modelled as a pressure term (red equation) describing cell contractility, and for the initial model we assumed a uniform distribution of contractility. Cell adhesions are distributed throughout the monolayer based on experimental observations (blue schematic), and at these points stresses within the monolayer are transmitted to the substrate to deform it (bottom two equations). Schematic produced in part with Biorender.com. **b,c** Total hydrogel displacement from experiments (same as Fig. S2g,h) were used to parameterise the model. **d,e** Based on this parameterisation the model can recapitulate the overall trend in experimental traction forces (same as Fig. 2e,f). **f,g** Model predictions on intercellular stresses at the radial midpoint of the layer (r=0.5) normalised to E=10kPa show changes as substrate stiffness increases. The model predicts intercellular forces change in a manner similar to the total traction forces as substrate stiffness increases. **h** Example time course projections of our laser ablation procedure for both micropatterns. A ring ROI with outer radius of 50µm and inner radius of 40µm was ablated. Merge shows overlay of 0s before and 60s after the ablation only to highlight deformations, where white highlights regions of no deformation. Scale bar = 20µm. **i,j** Mean recoil velocity vector plots for monolayers adhered on disks (**i**) and rings (**j**) on different substrate stiffnesses. Red dashed line outlines the adhesive surface and scale bars represent 3.5μm/min. **k,l** Quantification of maximum recoil velocity after ablation for monolayers adhered to disks and rings on different substrate stiffnesses. **m,n** Recoil velocity (same as Fig. 3k,l) and model predictions of radial midpoint intercellular stresses (same as Fig. 3f,g) have both been normalised to the value on the softest hydrogel (E=10kPa). The model is not able to mimic experimental observations. **o-r** Quantification of the total loss in traction forces (**o,p**) and strain energy (**q,r**) after ablations and once a new steady state is reached, for monolayers adhered to disks (**o,q**) and rings (**p,r**) on different substrate stiffnesses. Where **o’-r’** represent this value as a percentage of total traction forces (see Fig. 2e,f, **o’,p’**) and strain energy (see Fig. 2i,j, **q’,r’**). For all quantification graphs, experimental data is in black, with dots representing mean value, whiskers the S.E.M and grey line the line of best fit. Model outputs are red circles, where included.

As we observe a reduction in total adhesion coverage as substrate stiffness increases (Fig. S1d,e), we examined in our model the sensitivity of total traction forces to different levels of adhesion coverage (Supplemental Information). This revealed that traction forces remained relatively constant with different total levels of adhesion coverage above a 10% coverage threshold (Supplemental Information Table 2). As the coverage of adhesions is greater than 10% for all our conditions, we conclude that the reduction in adhesion coverage we observe does not drastically limit the traction stresses generated by the monolayer. Indeed, this analysis also shows that our results are insensitive to exact details of the underlying distribution of adhesions distributed throughout the layer. As the monolayer was stable at the timescales measured (Fig. S1k-m), we considered the system to be in a stationary steady state and the forces balanced. The model can be used to fit both the displacement data (Fig. 3b,c) and total traction (Fig. 3d,e) for each substrate stiffness, with the cell-generated contractile stress as the fit parameter. In fitting this data, we assume uniform contractility across the layer so that each cell is equally mechanically active. Where active matter models are used to describe traction force-type experimental systems the assumption of constant contractility is common^31–34,55^. Assuming that each cell across the layer had the same uniform contractility was sufficient to reproduce the observed traction force microscopy data, with small variations in this overall contractile activity on each stiffness. As with monolayer stress inference techniques, we then extended this analysis of traction forces to infer internal stress within the layer, akin to intercellular stress, which can be determined from the force balance equations. As a reference point, we plot the internal radial stress halfway across the radius (r=50μm; Fig. 3f,g). As might be intuitively expected, the model predicts internal stresses that broadly follow the same qualitative trends observed for total traction across both disks and rings. For the disks, we predict a significant monotonic decrease as substrate stiffness increases, with the greatest stresses being on the softer substrates (Fig. 3f). Whilst for the ring patterns, we see that the stress remains broadly constant across the stiffnesses (Fig. 3g). We next sought to examine experimentally whether the model’s internal stress predictions were accurate by considering both traction stresses using traction force microscopy and intercellular stresses inferred from laser ablation.

We bisected the epithelial monolayers with an annular region of interest (ROI) (Fig. 3h) to infer bulk intercellular stresses^25^, and analysed the recoil velocity (Fig. S3a,b). We observed that the greatest recoil was near the ablation site, and that vectors were orientated away from the cut highlighting that the monolayers were always under tension (Fig. 3i,j, Fig. S3c-f). Measuring the maximum recoil velocity half-life revealed a similar relaxation time of ∼20 ± 10s for both patterns and all stiffnesses. Assuming that the cells’ Young’s modulus is independent of substrate stiffness as previously reported^56^, this suggests that the viscoelastic properties of the monolayer were not altered by focal adhesion patterning or substrate stiffness (Fig. S3g-j). We then examined the maximum recoil velocity with respect to substrate stiffness for both patterns. For the disk pattern, the recoil velocity showed a weak decrease (Fig. 3k) qualitatively similar to the model prediction, but with the reduction in internal stress being more subtle than that predicted (Fig. 3m). On ring patterns, while the model predicts a constant recoil velocity, we observed a consistent decrease with increasing stiffness (Fig. 3l). This reduction, which is not intuitively expected or predicted by the model, suggests that a model with uniform mechanical activity does not describe the system fully and that additional factors must be at play.

Comparing traction forces before and after ablation also revealed a reduction, highlighting mechanical coupling of stresses within the monolayer to the tractions applied to the substrate (Fig3o,p). Extending this analysis to the strain energy, we also observed a reduction in strain energy after ablation on both patterns (Fig. 3q,r), with the exception for disks on the stiffest hydrogel where differences were negligible (Fig. 3q). As total traction and strain energy vary for the different patterns and substrate stiffnesses, we examined the percentage change to assess whether this reduction merely reflected the loss of cells in the ablation region (Fig. 3o’-r’). This revealed that the percentage loss of traction forces and strain energy (Fig. 3o’-r’), tended to be higher for rings than for disks. Taken together, this supports the assumption that there is stress transmission from cell-cell interactions which distributes forces into the substrate, and releasing intercellular stress through ablation leads to a reduction in traction stresses generated by the monolayer.

### Theoretical modelling suggests non-uniformity of contractility

To illustrate the incompatibility of a model with uniform contractility, where all cells generate equal active forces across the monolayer, with the observed recoil velocities we focused on the ring pattern which experimentally shows a constant total traction of ∼53.7MPa (Fig. 2f) but a reduction in recoil velocity as substrate stiffness increased (Fig. 3l). For uniform contractility (Fig. 4a) we used our model to examine how the constant contractility (*P*=*a*) would have to be altered to maintain a constant internal stress (represented in blue) or constant total traction force (represented in red) as substrate stiffness increased. To orientate, we plotted an illustrative contour of constant internal stress at r=0.5 (which corresponds to the ablation region, blue line) this enables us to identify a region below the contour (shaded in blue) which has a lower internal stress at r=0.5 (blue shaded region, Fig. 4a) as required. Likewise, we also plotted an illustrative contour of constant total traction (red line) and shaded the region above the contour which has greater traction forces (red shaded region, Fig. 4a). Note that with uniform contractility the shaded regions do not generate a practically relevant overlap (Fig. 4a), so that we cannot attain lower internal stress with constant traction, and a parameter sweep altering the contractility of the layer demonstrates that this is true for all physically relevant regimes (Supplemental information). This highlights that, as the system maintains a constant traction force (red contour) it does not encroach into the lower internal stress regime (blue shaded region). This would inevitably lead to a system whereby changes in traction forces would lead to similar changes in internal stress as previously predicted by monolayer stress inference techniques based on passive elastic models (Fig. 3f,g,^10,13,14,18^).

**Fig. 4.**
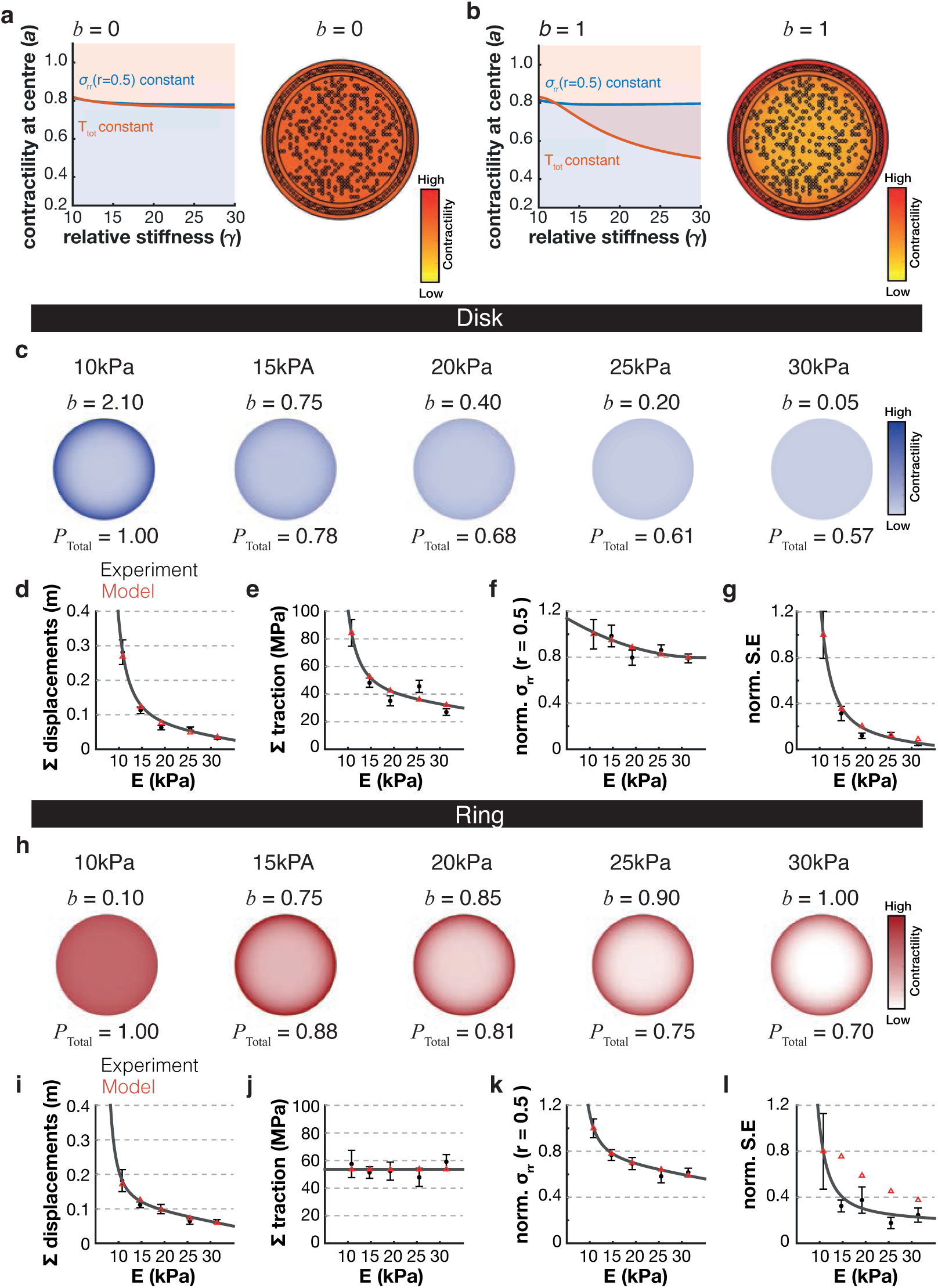
Non-uniform contractility reconciles experimental observations. **a,b** Phase plane plots of contractility in the centre of the monolayer against substrate stiffness for ring monolayers only. The examples showcase situations where the spatial distribution of contractility (*b*), represented with accompanying heatmaps, is either uniform (*b* =0, **a**) or non-uniform (*b* =1, **b**). In both, solid lines represent situations where either intercellular stress at the radial midpoint of the layer (r=0.5) is constant (blue lines) or total traction forces generated by the monolayer is constant (red line). The shaded regions represent either situations where the intercellular stress must be lower (blue shaded region) or total traction forces must be greater (red shaded area). Note that on uniform contractility there is no workable overlap of shaded regions meaning no workable parameters exist that allow constant traction forces but lower intercellular stresses as substrate stiffness increases as occurs experimentally. However, with non-uniform contractility an overlap region does exist, highlighting a large parameter space which could reconcile experimental observations. For **c-l** the model was parameterised to include non-uniform contractility for both disks (**c-g**) and rings (**h-l**). **c,h** Schematic representations of the total and spatial distribution of contractility parameters (*b* = spatial distribution of contractility; *P* = total contractility normalised to softest substrate) used for disk (**c**) and rings (**h**) on different substrate stiffnesses. **d,i** Total hydrogel displacement from experiments (same as Fig. S2g,h), **e,j** total traction forces (same as Fig. 2e,f), and **f,k** normalised intercellular stress at the radial midpoint of the layer (r=0.5) normalised to the softest substrate (same as Fig. 3m,n) were used to parameterise the model. **g,l** Normalised strain energy to the softest substrate was used as a final check to examine the power of the model to mimic all experimental observations. Note that for disks (**g**) the model can reproducibly mimic experimental observations but for rings (**l**) it mimics the trend but does not reproduce the actual magnitude of change. For all quantification graphs, experimental data is in black, with dots representing mean value, whiskers the S.E.M and grey line the line of best fit and model outputs are red triangles.

These phase plane plots do suggest a potential resolution by considering a non-uniform distribution of mechanical contractility within the layer. By increasing contractility at the edges, the system generates an overlap region where it is possible to maintain total tractions but reduce internal stress (Fig. 4b). In the examples given, to maintain a constant internal stress within the layer (blue line) traction forces must increase as the blue contour now encroaches the shaded orange region, and similarly to maintain constant total traction forces (red line) the internal stress must decrease as it now encroaches the blue shaded region (Fig. 4b). As the latter example is similar to our experimental observations, we explored whether parameterising the mechanical gradient could reconcile theoretical predictions with experimental observations.

We introduced a new contractility function that generates monotonic increasing contractility. Specifically, we take *P*(*r*) = *a*(1 + *br*^5^), as this enables analytical solutions in restricted cases and simplifies the numerical solution (see Supplemental Information). Here increasing *b* indicates more concentrated contractile activity at the edge (*b*=0 returns the model to uniform cell contraction). By parameterising the total contractility (*P*_Total_) of the monolayer and distribution of contractility (*b*), we were able to fit both tractions and relative changes in internal stress across the range of stiffnesses for both disk (Fig. 4c) and ring patterns (Fig. 4h). Namely we find that the model can reproduce a large decrease in traction and slight decrease in internal layer stress on disk patterns with increasing stiffness (Fig. 4d-f), while also reproducing nearly constant total tractions and large decrease in internal layer stress on the ring pattern as stiffness increases (Fig. 4i-k). Looking beyond the cell traction and recoil data, we considered the strain energy of the system. Non-uniform contractility within the model also reproduces the trends observed in the strain energy of both adhesive patterns (Fig. 4g,l), successfully capturing the observed reduction in strain energy for disks (Fig. 4g) but only qualitatively for ring patterns (Fig. 4l). However, considering that ring patterns have lower adhesion coverage than disks, a deviation in strain energy may be expected given the linear model of substrate resistance to deformation and the contribution of non-local effects from the thickness of the hydrogel (see Supplementary Information). Taken together, introducing non-uniform contractility into the model can recapitulate experimental observations.

Significantly, we find a qualitative difference in behaviour between disk and ring patterns for resolving experimental observations. For disk patterns the contractility must become more uniformly distributed, with more cells sharing the work as stiffness increases (Fig. 5a). However, on the ring pattern the opposite trend is required, with cells at the edge of the layer being required to be more contractile compared with the rest of the layer as stiffness increases, so that these edge cells do much of the work (Fig. 5a).

**Fig. 5.**
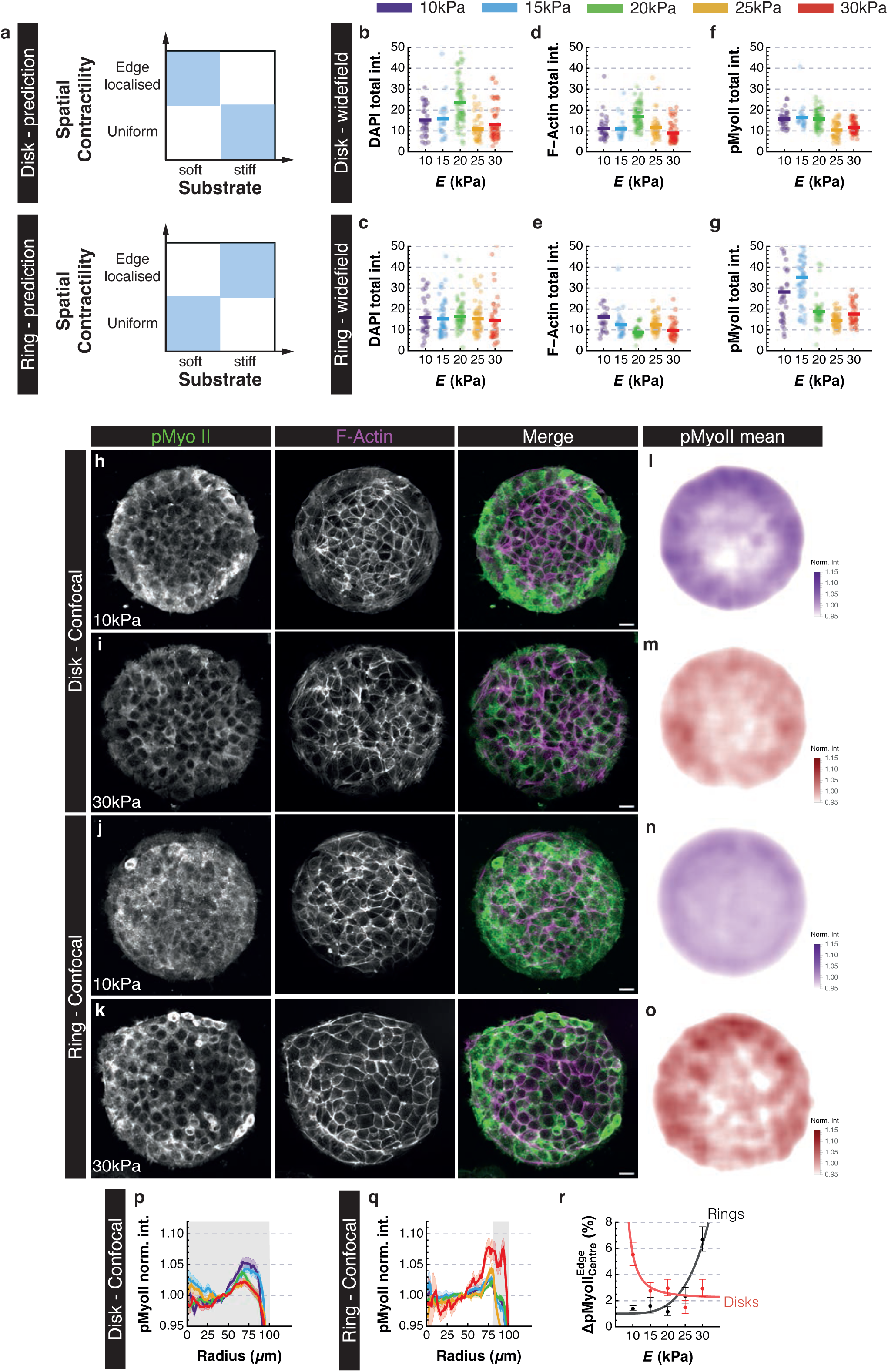
Contractility is non-uniform across micropatterned monolayers. **a** Schematic representation of predictions based on parameterisation of the model. For disks the model suggests that there needs to be a shift in contractility from focused to the edge on soft substrates to more uniformity on stiffer substrates. On rings, the model suggests the opposite needs to occur with contractility being more uniform on soft substrates and becoming more localised to the edge on stiff substrates. **b-g** Dot plots of total intensity from widefield images within each monolayer for DAPI (**b,c**), F-Actin (**d,e**) and pMyoII (**f,g**) adhered to disks (**b,d,f**) and rings (**c,e,g**). Each dot represents one monolayer, and the bars represent mean value for each population. **h-k** Example projections of total monolayer intensity taken from confocal sections of pMyoII and F-Actin adhered to disk (**h,i**) and ring (**j,k**) micropatterns on soft (**h,j**) and stiff (**i,k**) hydrogels. Scale bar = 20µm. **l-o** Heatmaps of mean total monolayer pMyoII intensity normalised to the ablation ROI (40-50µm) for monolayers adhered to disks (**l,m**) and rings (**n,o**) on soft (**l,n**) and stiff (**m,o**) hydrogels. **p,q** Quantification of radial mean total pMyoII intensity normalised to the ablation ROI (40-50µm) for monolayers adhered to disks (**p**) and rings (**q**) on different substrate stiffnesses. **r** Quantification of percentage difference in normalised total monolayer pMyoII intensity between edge (>60μm) and centre (<40μm) radii. Dots represent mean value, whiskers S.E.M and shaded line the line of best fit, with disks in red and rings in black. Note that the trend corroborates the model predictions (**a**).

### Active myosin and F-Actin are not uniformly distributed across monolayers

As our model suggested that non-uniformity of contractility could reconcile our experimental observations, we decided to examine contractility within the monolayers. We measured phosphorylated myosin light chain II (pMyoII) intensity across the monolayers as a proxy for contractility^20^, but also examined actin architecture which plays an essential role in cell mechanics^7^. As traction and intercellular stresses are dependent on myosin activity^20,57^, we initially wanted to examine if the total intensity of pMyoII altered depending on the pattern and substrate stiffness. To minimise experimental variation, we processed all independent samples in parallel to allow direct comparison in intensity using widefield imaging for speed. Measuring the intensity of DAPI (Fig. 5b,c), we observed consistent results with our cell count data (Fig. 1i,j). Examining F-Actin revealed that on disks there is a consistent level of F-Actin, that appears to be linked to cell number (Fig. 5d). However, on rings we observe a slight decrease in total F-Actin intensity as substrate stiffness increases (Fig. 5e). Next, we examined total intensity of pMyoII which for both patterns revealed a decrease as substrate stiffness increases, but more pronounced for rings (Fig. 5f,g). This agrees with our fitted model parameters which predicted a reduction in overall contractility as substrate stiffness increased (Fig. 4c,h).

To examine the spatial distribution of pMyoII, we imaged samples with improved optical resolution. Examining the total pMyoII intensity of z-sections across our monolayers we did observe spatial variation in pMyoII to varying degrees on our monolayers (Fig. 5h-k). For disk patterns, pMyoII intensity appeared more localised at the monolayer edge on soft hydrogels, becoming more uniform on stiffer hydrogels (Fig. 5h,i). For rings, the opposite trend in pMyoII intensity distribution was observed, with a more uniform distribution on soft hydrogels becoming more localised to the edge on stiffer hydrogels (Fig. 5j,k). As intensity was variable between samples, we normalised pixel intensity to the mean intensity within the ablation ROI (40-50μm) to quantify the spatial distribution of pMyoII (Fig. 5l-q).

This revealed that no condition had a uniform localisation, with radial variation always present (Fig. 5l-q), however, there was a redistribution of pMyoII for both patterns as substrate stiffness increases. On disks there was a reduction in pMyoII intensity outside the ablation ROI (Fig. 5p), whilst on rings an increase was observed (Fig. 5q). Measuring the percentage difference in pMyoII intensity between the edge and central regions (Fig. 5r) revealed a behaviour similar to that predicted by our model (Fig. 5a); as substrate stiffness increased, on disk patterns there is a trend for pMyoII to become more uniform but on rings patterns there is a trend to become more localised at the edge.

As there are minor differences in cell height on ring patterns (Fig. 1l), we wanted to determine if cell height was an issue in this analysis. We therefore examined pMyoII localisation across the cell height which revealed that it was predominantly localised basally (Fig. S4a-c). Consequently, we examined actomyosin distribution across the monolayers focusing only on the basal layer (Fig. S4d-l). Examining pMyoII we observed similar changes in distribution across the basal layer as we did the whole monolayer (Fig. S4j-l), suggesting that cell height wasn’t confounding our analysis. Interestingly, this analysis revealed drastic changes in the distribution of the actin cytoskeleton between patterns (Fig. S4d-i) which was masked when examining total intensity, as F-actin tended to be more localised along cell junctions (Fig. S4a-c). On disk patterns, cells at the periphery extended lamella outwardly leading to a dense band of F-Actin at the edge (Fig. S4d,f,h). This was followed by a reduction in F-Actin behind this band, as cell nuclei aligned close to the edge, with the more centrally located cells exhibiting clear fibres that spanned the cell width and were predominantly located near active focal adhesions (Fig. S4m, m’). On ring patterns, an F-Actin bundle spanning the monolayer perimeter was observed, with a reduction in F-Actin along the laminin ring, and another accumulation at the inner ring diameter but not as a defined actin bundle (Fig. S4e,g,i). The architecture of the actin cytoskeleton is critical for generating^58^ and transmitting^17^ stresses, and the differences between ring and disk patterns highlights how altering focal adhesion distribution drastically alters the mechanical equilibrium of the tissue.

Overall, reconciling the model and experimental observations, we find that the relationship between traction and intercellular stress is not universal but changes in response to both the distribution of focal adhesions and compliance of the substrate. This is due to cellular contractility adapting, to generate a tissue-wide force balance.

## Discussion

The control of a tissue’s mechanical equilibrium is essential for the regulation of a wide array of physiological processes such as morphogenesis^1^ or wound healing^3^, and when disturbed can lead to disease progression^59^. Here we investigated the relationship between applied surface traction and intercellular stresses specifically focusing on the role of focal adhesions in the mechanical equilibrium of epithelial monolayers. We find that, for ring patterns where focal adhesion distribution is non-uniform, epithelial monolayer stresses do not conform to the intuition that the relationship between traction and intercellular stresses is linear (Fig. 3m,n). Instead, while traction forces do not change upon increased substrate stiffness (Fig. 2f,h), intercellular tension as measured by laser ablation dramatically drops (Fig. 3l). To reconcile experimental observations with the monolayers being at force-balance, we proposed and confirmed that spatial patterns in contractility emerge (Fig. 4, 5).

What is the origin of this apparently counter-intuitive behaviour? In comparison with single cell studies, for large clusters of cells, forces can be distributed differently between the cells and substrate according to the state of cell-cell junctions^55,60^ and the distribution of cell-ECM attachments^16^. In particular, attachment to the ECM has been shown to effectively decrease the length-scale of stress transmission^30^, so differences in substrate stiffness can potentially be transmitted to cell-cell junctions over longer distances if cells are not all anchored to the matrix. Conversely, as with the disk patterns where focal adhesions are widely distributed (Fig. 1g), often near cell-cell junctions (Fig 1e) and associated to the contractile acto-myosin network (Fig. S4m), cell-generated forces are most likely transmitted locally to the substrate. In this situation the mechanical properties of the substrate become important for stress transmission across the tissue, as has previously been reported for fibroblasts^61^. Indeed, we found that on disk patterns the monolayer was less able to deform stiffer hydrogels (Fig. 2c), and this was accompanied by a decrease in total traction (Fig. 2e). This is associated with a reduction in the overall total contractility of the layer on stiffer gels (lower pMyoll total intensity, Fig. 5f) as predicted by our theoretical model (Fig. 4c). On soft conditions however, mechanical stress propagates further across the monolayer as cells are more able to deform the substrate locally (Fig. S2e), and higher total contractility is observed (Fig. 5f). This is accompanied with contractile activity differentially localised most strongly to the edge of the layer (fig. 5l,p) where the highest deformations were measured (Fig. S2e).

In contrast, for ring patterns on increasing substrate stiffness, traction stress is relatively stable (Fig. 2f,h) whilst intercellular tension decreases (Fig. 3l), highlighting that different mechanical states are reached. For rings, it is only the cells at the edge of the monolayer that can form focal adhesions and directly experience gel deformability (Fig. 1h). The bulk of the monolayer can now only contribute predominantly to stress distribution through cell-cell junctions. Similar to the disk patterns, there is still a decrease in deformation on stiffer gels (Fig. 2d) and a corresponding decrease in total contractile activity of the monolayer (lower pMyoll total intensity, Fig. 5g). However, the non-uniform adhesion distribution generates a marked shift in the collective behaviour of the tissue.

With increasing stiffness, the mechanical activity becomes progressively more localised to the edge of the monolayer on stiffer gels (Fig. 5r), thus generating lower internal stress as observed and predicted (Fig. 4k). The stronger resistance to deformation of stiffer gels has an anchoring effect on the outer ring, but now the balance of resistive forces with the unadhered cells adjacent to these ring cells is altered. These results emphasise that the processes contributing to force transmission throughout a monolayer cannot be isolated to a single cell without considering their collective effects. This is highlighted by changes in F-Actin architecture across the monolayer between disk and ring patterns (Fig. S4).

Changes in the spatial distribution of actomyosin contractility underpins the response to differential adhesion on our ring and disk patterns (Fig. 4, 5). Spatial distribution of contractility has been observed in single cells^62,63^ linked to focal adhesion signalling^62^, however for epithelial monolayers, whilst radial patterns of focal adhesion distribution have been reported^64^, to our knowledge this has not been linked to non-uniform patterns of contractility. Spatial variation in myosin have been observed *in vivo*, but these are normally examined in the context of morphogenesis^65^ or generating cell-population boundaries^66^ and not linked to cell-ECM attachments. We recently observed in the *Drosophila* abdominal epidermis that a tissue-wide spatial pattern of myosin arises during the pupal stages as the ECM is degraded^27^ but whether this is important for the tissue’s mechanical equilibrium is not known. However, the emergence of spatial patterns of contractility in our monolayers has important implications on the application of force inference techniques. To examine intercellular stresses of a monolayer, inference methods such as monolayer force microscopy (MSM)^10,11,18^ or Bayesian inversion stress microscopy (BISM^12^) have been developed. These techniques infer intercellular stresses based on traction forces measured and can estimate spatial variation in intercellular stresses in a non-destructive manner, unlike laser ablation, allowing long-term analysis to be made. The implementation of these techniques often assumes a passive elastic model linking traction and intercellular stresses^10,11,13,17,67^. For MDCK cell-doublets, where focal adhesions have been shown to disassemble along cell-cell contacts^68^, the direct balance of traction forces with cell-cell adhesions potentially holds^14^. However, even in these simple cell-doublet systems the distribution of focal adhesions can drastically alter the propagation of forces. For instance, cardiomyocyte doublets seeded or cultured on 90kPa substrates generated focal adhesions at the site of cell-cell junctions, impairing mechanical coupling and force propagation^15^. This highlights the importance of examining the distribution of focal adhesions when inferring stress transmission. Moreover, on our 200μm diameter disk micropatterns, we observed spatial variation in pFAK with some regions having higher intensity than others (see Fig 1c). Considering a larger tissue, the distribution of focal adhesions could vary drastically even on a uniform substrate, altering the local mechanical equilibrium in that region of a tissue. This would impact whether local intercellular stresses were positively correlated with traction stress or not. Indeed, analysis of epithelial monolayers showed a “rugged landscape” of traction forces^10^, and spatial patterns of ECM composition and properties have been observed in developmental and pathological situations^41^. Therefore, when employing techniques such as MSM and BISM it is important to not only examine the variation in local distribution of focal adhesions, but also spatial variation in contractility to confidently infer intercellular stresses as a function of the mechanical activity of the tissue as a whole.

## Methods

### Cell culture

EpH4 cells with Cas9 inserted into the Rosa26 locus (kind gift from high throughput screening STP, Francis Crick Institute) were cultured in 4.5g l^-1^ Dulbecco’s modified Eagle medium (Gibco, Thermo Fisher Scientific) containing 10% foetal bovine serum (Sigma-Aldrich, Merck) and 1% penicillin-streptavidin (Gibco, Thermo Fisher Scientific). For seeding on hydrogels, 180,000 cells in 2ml of culture medium were centrifuged at 80 r.c.f for 5 minutes. Cells were cultured for 2 hours and then extensively washed in PBS to remove non-adherent cells and 2ml of fresh media was added. Cells were incubated for 48 hours, to proliferate and cover the micropatterns. For live-imaging experiments, cells were washed in PBS and then labelled for 15 minutes with 1.5μl CellMask Green (Thermo Fisher Scientific), diluted and vortexed in 1.5ml FluoroBrite DMEM (Gibco, Thermo Fisher Scientific) containing 1% penicillin-streptavidin but no FBS. Media was then replaced with 1.5ml FluoroBrite DMEM media as described, ready for imaging.

### Micropatterning of methacrylated hyaluronic acid hydrogels

3g of methacrylated hyaluronic acid (MeHA) was synthesised as previously described^43^, with a methacrylate modification of hyaluronic acid monomer between 76 – 79% as characterised through ^1^H NMR (Fig. S1a). MeHA hydrogels were formed at a concentration of 3 wt% and crosslinked with increasing amounts of dithiothreitol (DTT) to modulate substrate rigidity. Specifically, hydrogels were formed within a 250μm high silicon gasket with a 16mm inner diameter and outer diameter of 18mm, sandwiched between a 19mm micropatterned coverslip and a silanized glass surface. Hydrogels were incubated with 5% BSA (Tocris Bioscience) in PBS for 2 hours before seeding cells.

Micropatterned coverslips were generated by plasma-activating 18mm coverslips and coating in 0.1mg ml^-1^ poly-L-lysine poly(ethylene glycol) (PLL(20) – g[3.5]-PEG(2), SuSoS) suspended in 10mM HEPES buffer at pH7.4. To pattern the coverslips, a quartz photomask (Compugraphics) and UVO-cleaner (Jelight) were used to remove PLL-g-PEG from the surface of the coverslip. Patterned coverslips were then incubated for 30 minutes with laminin diluted in 100mM sodium bicarbonate buffer at pH7.4. Laminin consisted of either 20μg ml^-1^ laminin (Sigma-Aldrich, Merck) solution or 20μg ml^-1^ 2:1 mix of laminin and laminin-labelled with alexa633 for live-imaging and immunostaining, respectively. To label laminin, 500μl of 1mg ml^-1^ laminin solution was injected into a Slide-A-Lyzer (Thermo Fisher Scientific) and dialysed two times in 1L of PBS at 4°C and stirred at a low shear rate for 24 hours. The dialysed laminin was then labelled with Alexa633 following the manufacturers protocol (Thermo Fisher Scientific). Micropatterns were either repeating patterns of disks with a diameter of 200μm, or rings with an outer diameter of 200μm and inner diameter of 160μm.

For silanized glass surfaces, either No. 1.5 glass bottom 35mm μ-dishes high (Ibidi) or 20mm diameter uncoated 6-well plates (Mattek) were employed for live-imaging or immunostainings, respectively. Glass surfaces were plasma-activated and silanized with (3-aminopropyl) trimethoxysilane (VWR) and 5% glutaraldehyde (VWR).

### Atomic force microscopy

Gels were generated as described but using non-patterned, passivated coverslips and No. 1.5 50mm dish with 30mm uncoated glass diameter (Mattek). Force curves were measured on a JPK Nanowizard 4 using a CP-qp-CONT-Au-B-5 cantilever probe (sQube, NanoAndMore GMBH). Measurements were taken in four regions per hydrogel (two at the periphery and two central) with 50 indentations over a 100μm x 100μm grid. The number of hydrogels quantified for each DTT concentration was 7 for 6.30 to 15.75mM and 6 for 18.9mM. Force curves were made with an extend speed of 2μm s^-1^ and a setpoint (8 – 25nN, depending on substrate stiffness) set to achieve a maximum indentation depth of ∼500nm. The young’s modulus was calculated by fitting a Hertz model using the AFM_Youngs_modulus_fit programme developed by Dr Stefania Marcotti (available on GitHub: https://github.com/INSIGNEO/AFM_Youngs_modulus_fit), at different indentation depths (Fig. S1b). A standard indentation depth of 250nm across all hydrogel conditions was used to calculate the Young’s modulus, as it is within the linear phase of the modulus versus indentation depth for all hydrogel conditions and is 1/1000 of the hydrogel thickness.

### Traction force microscopy

Crimson labelled 0.2μm carboxylate modified FluoSpheres (Thermo Fisher Scientific) were mixed with MeHA hydrogel precursor solution with a 1:40 ratio prior to crosslinking with DTT. Images were acquired as detailed below. After live-imaging, cells were treated with 10x trypsin-EDTA, no-phenol red (Gibco) until cells detach from the hydrogels and a final reference frame acquired.

### Laser ablation

Images were acquired on a Zeiss LSM 780 system with an LD C-Apochromat 40x NA 1.1 water objective and 0.8 zoom, so that micropatterns could be acquired in a single 512 x 512 pixel tile. Z-stacks were acquired by measuring excitation by 480nm and 633nm lasers simultaneously for z-slices every 1μm, spaced between 5 - 6μm below the hydrogel surface and 10 - 12μm above. Z-stacks were acquired every 10 seconds for one minute before a laser ablation and five minutes afterwards. An annulus ROI was generated using the freehand tool and had an internal diameter of 80μm (154 pixels) – approximately 4 – 6 cells in diameter – and external diameter of 100μm (193 pixels) – leaving 3 – 4 cells at the periphery. Ablations were made in a z-plane positioned in the middle of the epithelial layer (3 - 6μm from the hydrogel surface) with a Coherent Chameleon NIR tuneable laser set at 780nm and 40% - 60% power output and lasted ∼6.7ms.

### Immunostaining

Samples were fixed in 4%-paraformaldehyde (TAAB) for 15 minutes and permeabilised with 0.3% Triton X-100 for 5 minutes, covered from light. Samples were then blocked with 10% neonatal goat serum (Generon) for an hour at room temperature and incubated with primary antibodies diluted in 10% neonatal goat serum at 4°C overnight. Samples were then incubated in secondary antibodies, Phalloidin-Alexa568 (Thermo Fisher Scientific, #A12380: dilution 1:400) and DAPI (1000x stock solution; Sigma) for 1 hour at room temperature. Samples were washed and stored in 0.1% Triton X-100. For batch processing of all samples to examine total pMyoII levels, images were acquired on a Zeiss Celldiscoverer 7 widefield system with a Zeiss Plan-Apochromat 20x NA 0.7 Autocorr objective with 1x zoom and 2240 x 2254 pixels and a x-y resolution 0.1747602 for a single plane. To examine spatial localisation of proteins of interest, samples were imaged on a Nikon CSU-W1 spinning disk with an Apo LWD 40x WI λS DIC N2 NA 1.15 objective with 1200 x 1200 pixels and a x-y resolution of 0.2731993μm. Z-slices were acquired 0.4μm apart for each channel sequentially. The following antibodies were used, mouse anti-phospho-Myosin Light Chain 2 (Ser19) (Cell Signalling Technology, #3675: dilution 1:200), rabbit anti-phospho-FAK (Tyr397) (Thermo Fisher Scientific, 44-624G: dilution 1:400), mouse anti-E-Cadherin (BD Transduction #610182: dilution 1:400), Goat anti-mouse IgG H&L Alexa 405 (Abcam, #175660: dilution 1:400), and Goat anti-rabbit IgG H&L Alexa 488 (Thermo Fisher Scientific, #A-11008: dilution 1:400). F-Actin was stained with Alexa Fluor 568 Phalloidin (Thermo Fisher Scientific, #A-12380) along with the secondary antibodies.

### Data analysis

All images were processed and analysed in Fiji, Mathworks MATLAB and Wolfram Mathematica.

### Traction force microscopy pipeline

Images were acquired as detailed above. After adding trypsin to detach cells, there is slight stage movement in the in x, y and z directions. Therefore, all movies were registered with the trypsin frame as a reference. For z registration, the surface of the hydrogel was calculated by examining the total intensity across the image in 24×24 pixel blocks for each z-slice, with the surface for each block set by the last slice with a signal. To project on a pixel-by-pixel basis, a Gaussian Filter was employed across all blocks and then rounded, to minimise any jumps in z-plane. A maximum projection of the beads was then calculated for both the movie and trypsin reference frame from the surface to the same number of z-slices below the surface set by the least number of frames available, normally 3 - 4 slices. Once z-registration is complete the trypsin frame was added as the final frame of the movie and registered in Fiji with the StackReg macro, using the last frame as the reference. Bead displacement, traction forces and strain energy were calculated using the TFM package developed by Gaudenz Danuser’s group^53^, using a 31×31 pixel template size and L-parameterisation of 0.00000001.

### Laser ablation analysis

Initially a projection of the cell mask was generated by identifying the ablation plane and calculating a maximum projection between two slices above and below this frame. The ablation plane was identified by measuring the loss of intensity within the ablation ROI for each frame after the ablation and selecting the frame with the biggest loss. To measure the recoil velocity, the Mathematica ImageFeatureTrack function was employed with specifying features in every pixel and filtering pixels where no motion was detected. To measure maximum recoil velocity for each movie, the recoil velocity value for the 95^th^ percentile across the entire field of view was taken (see Fig. S3).

### Cell morphology quantification

To measure cell elongation and area, projections generated for analysing the recoil velocity were segmented through Skeletor^27^ and manually corrected in Fiji. Using a watershed algorithm in Mathematica, cell area and elongation (1-width/length) were measured. To measure cell height, all immunostaining experiments were combined, and an orthogonal projection made using the Fiji macro with a box ROI 20 x 850 pixels centred on the monolayer centroid in both the x and y direction. In Mathematica, cell height was measured by combining F-Actin intensity with either pMyoII or E-Cad intensity for the orthogonal projections, depending on the experiments, and binarizing based on pixels whose intensity was greater than the mean plus four standard deviations of the entire projected image. A FillingTransform and DeleteSmallComponents function were applied to clean up the binarization so only the monolayer remained.

### Membrane intensity changes

To measure the movement of the epithelium in our movies before the ablation to determine their dynamics, the frame immediately before the ablation was subtracted from the initial frame and the remaining pixel values summed and normalised to the original summed pixel values. A value close to zero highlights very little movement within the epithelium.

### Basal projection algorithm

To quantify just the basal layer of the monolayers we used Laminin labelling to identify the surface of the hydrogel. We selected a z-slice for each pixel that corresponded to the highest laminin intensity, after noise-filtering the z-intensity profile with a LowpassFilter. We then generated a contour map by smoothening the pixel values with a GaussianFilter, before rounding to a whole number to correspond to a z-slice. Based on this contour map, we then did a maximum projection over three z-slices, one above and one below our contour map value.

### pFAK segmentation

To quantify adhesion coverage and size, pFAK images were binarized based on pixel intensity value in Mathematica. Due to variable background intensity across the field of view, a MedianFilter was applied with a 61 pixel-by-pixel sliding window and subtracted from the original image. A background threshold value was then calculated by measuring the mean intensity plus two standard deviations of pixels outside the micropatterned region which was determined based on the laminin micropattern channel. Any pixel which had a higher intensity value than the background threshold was binarized. To de-noise the binary image, isolated binarized pixels were removed using the DeleteSmallComponents function with a value set to 1.

### Statistical analysis

All statistics were performed in Mathematica using the LocationEquivalenceTest function which tests the null hypothesis that populations are equal choosing the most powerful test that applies to the data. The most common test applied was the Kruskal Wallis test as often one condition would fail normality tests. If the LocationEquivalenceTest was significant a post-test Mann-Whitney was performed comparing the value on the softest substrate to all other substrate stiffnesses. For trend fitting, the LinearModelFit function was used on the mean values for each population with a confidence interval of 95%. The model with the greatest R^2^-value was selected.

For all statistical values used in the analysis, see Tables S1&2 in Supplementary Information.

### Theoretical modelling

We describe the theoretical and computational modelling briefly here, for a more detailed description see the additional supplementary information.

### Active matter model for the cell layer

An active matter model was used to predict the deformations and stresses in the cell layer. This widely adopted approach incorporates an active cell-generated stress additively into the constitutive relation of the cell material, see^36,37,39,54^. As such we assumed a constitutive relation such that the stress ***s* = *s*_passive elastic_ + *s*_active_,** as Fig. 3a specifically in tensor notation this gives:

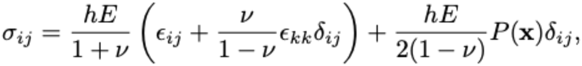

where we use the summation convention and introduce the kronecker delta. The active contractility P(*x*) is assumed to act isotropically. The deformations are then determined from the force balance

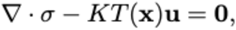

Where *K* quantifies substrate resistance to deformation and *T*(x) is an indicator function that take the value of 1 where the cell is adhered and 0 otherwise. The contractility function is assumed to be constant throughout the layer *P*(x)=*P*_a_, for Fig. 3, with the addition of a monotonic increasing contractility profile *P*(x)=*a*(1+*b*r^5^) for Fig. 4. The polynomial contractility profile is selected to enable analytical phase plane analysis. Where adhesion is uniform analytical solutions of the force balance equation may be obtained by standard techniques as the resulting equations are of Bessel type and solutions may be obtained by variation of parameters. Where non-uniform adhesion is observed numerical solutions of the force balance equation are obtained using finite element methods within the MATLAB PDE Toolbox.

### Adhesion geometry

For uniform adhesion the indicator function is either taken as unity throughout for the disk geometry or as unity in the ring region. Where adhesion is informed by the pFAK data (Fig. 1) and modelled as non-uniform we take the resulting adhesion geometry is as in Fig. 1h. Specifically, we consider two thin annuli of thickness *r_t_*=0.06 at the edges of the micropatterned ring corresponding to the regions with highest intensity pFAK. Additionally, we set a distribution of adhered spots in the internal region and between the two annuli as the pFAK intensity is non-zero in these regions; spots between the two annuli have a radius of *r_s_*=0.0167*r_0_* and in the internal region have radius *r_s_*=0.018*r_0_*. In these adhered regions T(x)=1 and elsewhere we set T(x)=0.

## Acknowledgements

We would like to thank the Ok-Ryul Song and Michael Howell from the High-throughput screening STP at the Francis Crick Institute for sharing the EpH4-Cas9 cells. We are grateful to Matt Renshaw from the Crick CALM STP, and Ravi Desai and Christina Dix from the Crick Making STP for their assistance. We thank the bioAFM and bioImaging facilities within FBMH at the University of Manchester. We are grateful to Stefania Marcotti, Richard Thorogood, Nigel Hodson, Olivia Courbot and Alberto Elosegui-Artola for advice on the analysis of AFM data. We thank Guillaume Charras and Alberto Elosegui-Artola for comments on the manuscript. JRD was supported by a Wellcome Trust Sir Henry Wellcome fellowship (201358/Z/16/Z) and University of Manchester Dean’s Prize. JSW was supported by postdoctoral EPSRC funding (EP/W522302/1). This work was supported by a Wellcome Trust Investigator award (107885/Z/15/Z) to NT. Work in the Tapon lab is supported by the Francis Crick Institute, which receives its core funding from Cancer Research UK (CC2138), the UK Medical Research Council (CC2138), and the Wellcome Trust (CC2138). For Open Access, the authors have applied a CC BY public copyright licence to any Author Accepted Manuscript version arising from this submission.

---

**Fig. S1.**
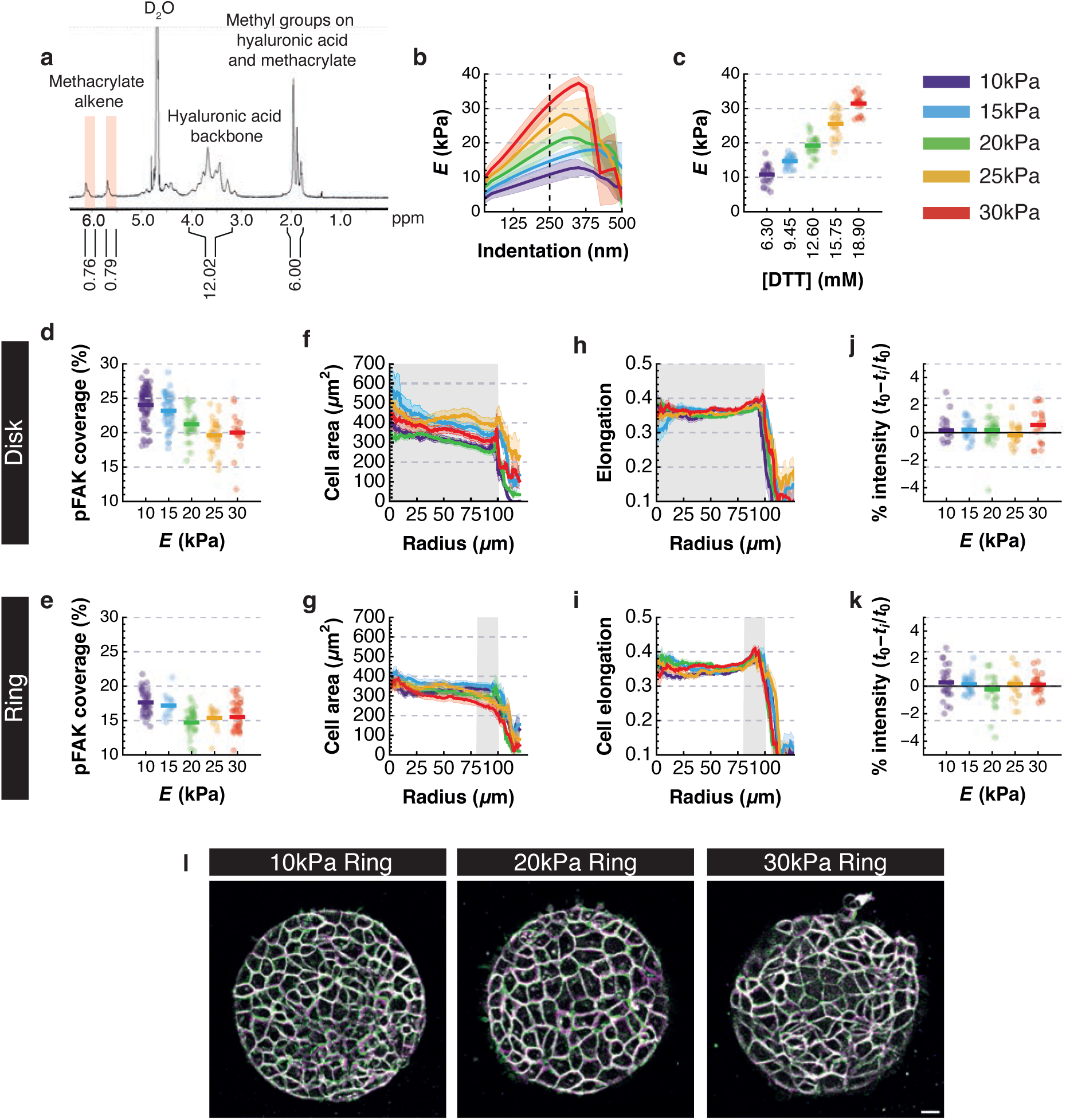
Characterisation of micropatterned MeHA on EpH4 cells. **a**^1^H NMR spectra of methacrylated hyaluronic acid (MeHA). **b** Young’s moduli at different indentation depths on 3%(w/v) MeHA hydrogels cross-linked with different DTT concentrations. Dashed line is at 250nm which was used to define the Young’s moduli for each DTT concentration. **c** Dot plot of Young’s moduli for each concentration of DTT. Each dot is the average value from 50 force-curves taken for a 100μm x 100μm region. **d,e** Dot plot of total pFAK coverage within each monolayer, bars highlight mean value for each population. **f,g** Quantification of mean cell area radially across micropatterns. **h,i** Quantification of mean cell elongation radially across micropatterns. For **f-i** thick line are mean values and ribbons are S.E.M, grey shaded region represents adhesive surface. **j,k** Dot plot of % intensity change after subtracting the cell outline projection from the frame before an annular ablation from the initial frame 1 minute before. Lines represent mean value for each population. **l** Cell outline projections five minutes apart overlayed with the initial frame in green and last frame in Magenta. White represents pixels where there is no change in intensity. All N numbers and statstics included in Table S2 in Supplementary Information. All scale bars = 20μm.

**Fig. S2.**
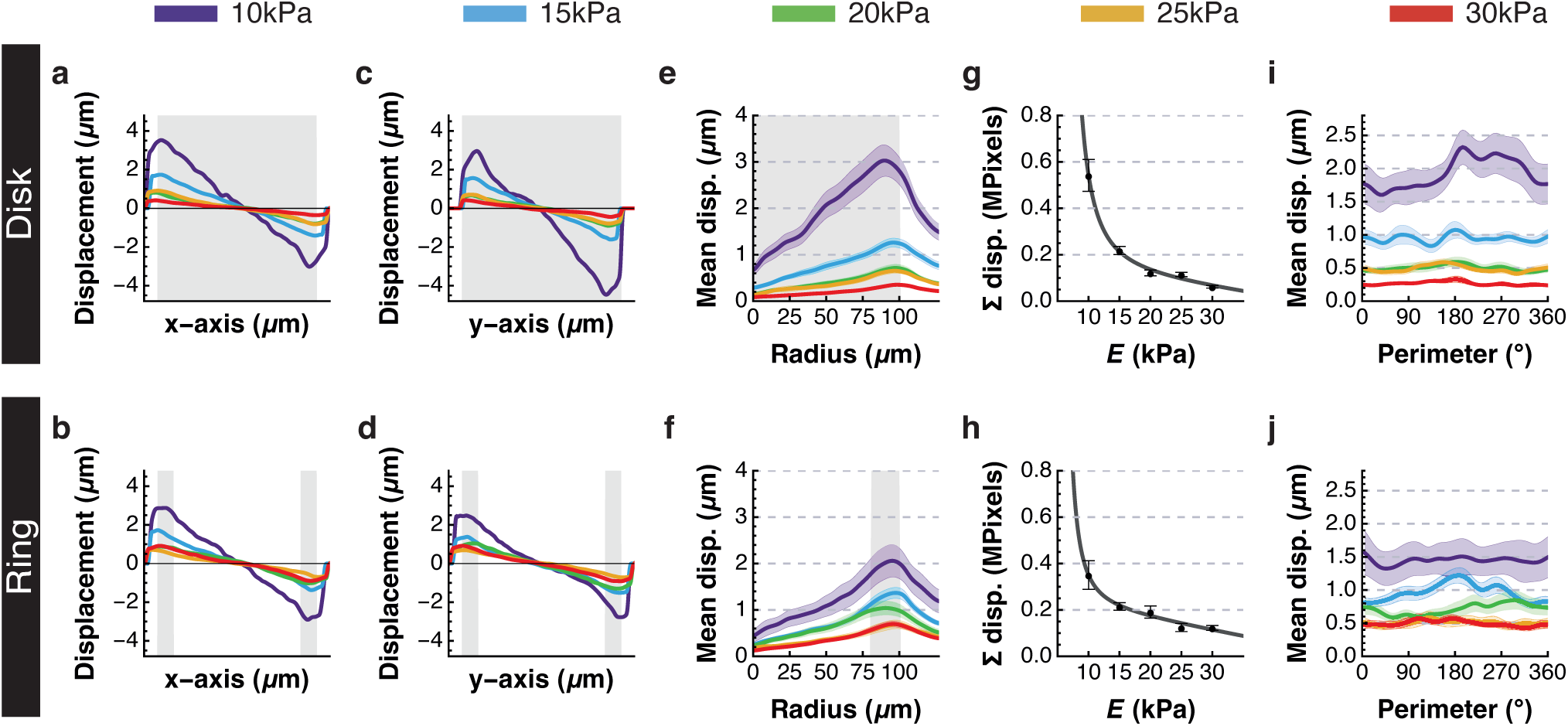
Displacements of microbeads embedded within MeHA hydrogels by EpH4 monolayers. **a-d** Quantification of mean displacement along the x-axis (**a,b**) and y-axis (**c,d**). Grey region represents adhesive surface. **e,f** Quantification of mean displacement radially across micropatterns, where thick lines are mean values, ribbons are S.E.M, and grey shaded region represents adhesive surface. **g,h** Quantification of total bead displacements generated by monolayers adhered to disks and rings on different substrate stiffnesses. Dots represent mean values, whiskers are S.E.M, and grey line is line of best fit. **i,j** Quantification of the mean bead displacement around the perimeter of monolayers, with 0° along the horizontal axis, where thick lines are mean values, ribbons are S.E.M, and grey shaded region represents adhesive surface.

**Fig. S3.**
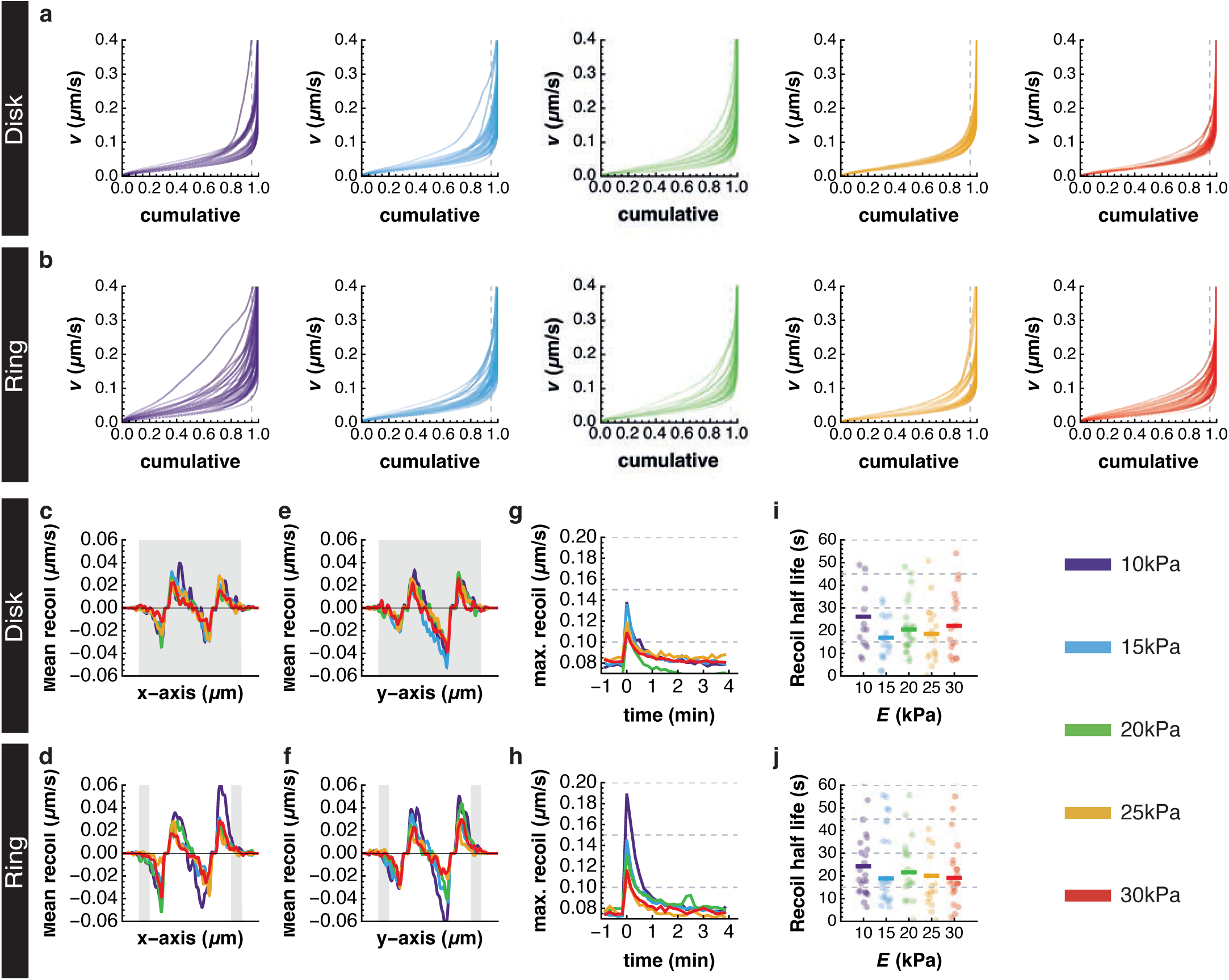
Recoil velocity dynamics and correlations after laser ablation. **a,b** Cumulative plots of all recoil velocity values for each movie on both micropatterns. To calculate the maximum recoil value the 95^th^ percentile (grey dashed line) value was used, which was always located near the edge of the ablation site. **c-f** Quantification of mean recoil velocity along the x-axis (**c,d**) and y-axis (**e,f**). Grey region represents adhesive surface. **g,h** Quantification of the mean maximum recoil velocity over the time course of our movies for both micropatterns. **i,j** Dot plot of recoil half-life calculated from Fig. S3g,h. Each dot represents a movie, and the line represents population mean value.

**Fig. S4.**
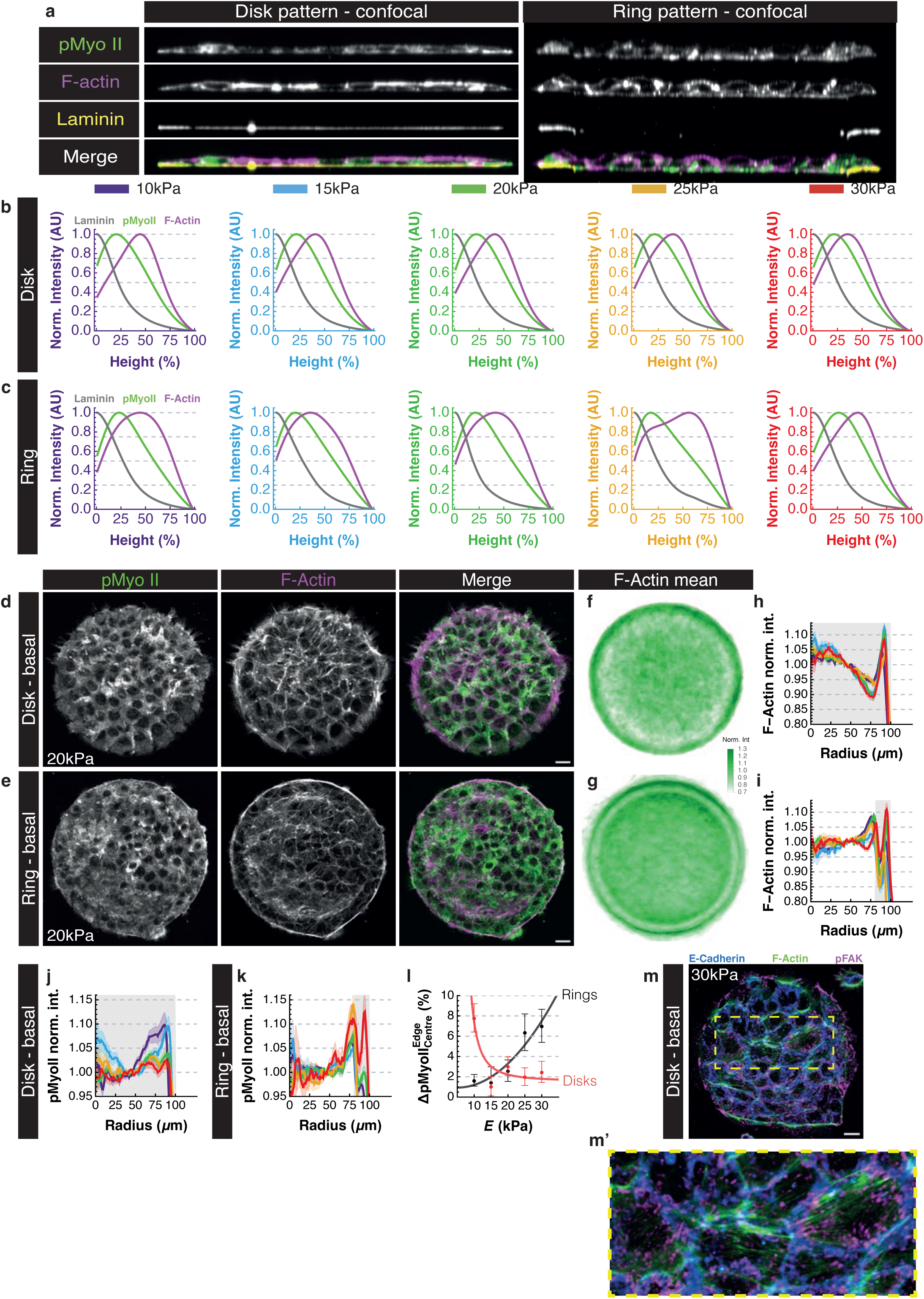
pMyoII localisation within monolayers. **a** Example orthogonal projections through the middle of a monolayer of pMyoII, F-Actin and Laminin on disk (left) and ring (right) micropatterns. **b,c** Quantification of mean orthogonal intensity of laminin, pMyoII and F-Actin normalised to the maximum value for the entire orthogonal projection (see **a**). Note that pMyoII maximum intensity is always near the laminin intensity highlighting basal accumulation of pMyoII, whilst F-actin peak is close to 50% of cell height where cell-cell junctions are located. **d,e** Example projections of basal monolayer intensity taken from confocal sections of pMyoII and F-Actin adhered to disk (**d**) and ring (**e**) micropatterns on 20kPa hydrogels. **f,g** Heatmaps of mean F-Actin intensity from basal projections normalised to the ablation ROI (40-50µm) for monolayers adhered to disks (**f**) and rings (**g**) on 20kPa hydrogels. **h,i** Quantification of radial mean F-Actin intensity from basal projections normalised to the ablation ROI (40-50μm) for monolayers adhered to disks (**h**) and rings (**i**) on different substrate stiffnesses. **j,k** Quantification of radial mean pMyoII intensity from basal projections normalised to the ablation ROI (40-50μm) for monolayers adhered to disks (**j**) and rings (**k**) on different substrate stiffnesses. For **h-k**, thick lines are mean values, ribbons are S.E.M, and grey shaded region represents adhesive surface. **l** Quantification of percentage difference in normalised pMyoII intensity from basal projections between edge (>60μm) and centre (<40μm) radii. Dots represent mean value, whiskers S.E.M and shaded line the line of best fit, with disks in red and rings in black. Note a similar trend to total projections seen in Fig. 5r). **m** Example basal projection of pFAK, F-Actin and E-Cadherin on disk micropattern with expanded insert (**m’**) to highlight F-Actin fibres that span epithelial cells, mainly between pFAK labelled adhesions. Scale bar = 20μm.

## References

1. Clarke, D. N. & Martin, A. C. Actin-based force generation and cell adhesion in tissue morphogenesis. Current Biology 31, R667–R680 (2021).

2. Pandya, P., Orgaz, J. L. & Sanz-Moreno, V. Actomyosin contractility and collective migration: may the force be with you. Curr Opin Cell Biol 48, 87–96 (2017).

3. Lim, S. E., Vicente-Munuera, P. & Mao, Y. Forced back into shape: Mechanics of epithelial wound repair. Curr Opin Cell Biol 87, 102324 (2024).

4. Mao, Y. & Baum, B. Tug of war-The influence of opposing physical forces on epithelial cell morphology. Dev Biol 401, 92–102 (2015).

5. Romani, P., Valcarcel-Jimenez, L., Frezza, C. & Dupont, S. Crosstalk between mechanotransduction and metabolism. Nat Rev Mol Cell Biol (2020) doi:10.1038/s41580-020-00306-w.

6. Dupont, S. & Wickström, S. A. Mechanical regulation of chromatin and transcription. Nature Reviews Genetics vol. 23 624–643 Preprint at 10.1038/s41576-022-00493-6 (2022).

7. Sun, X. & Alushin, G. M. Cellular force-sensing through actin filaments. FEBS Journal 290, 2576–2589 (2023).

8. Gómez-González, M., Latorre, E., Arroyo, M. & Trepat, X. Measuring mechanical stress in living tissues. Nature Reviews Physics 2, 300–317 (2020).

9. Lekka, M., Gnanachandran, K., Kubiak, A., Zieliński, T. & Zemła, J. Traction force microscopy – Measuring the forces exerted by cells. Micron 150, (2021).

10. Tambe, D. T. et al. Collective cell guidance by cooperative intercellular forces. Nat Mater 10, 469–75 (2011).

11. Tambe, D. T. et al. Monolayer Stress Microscopy: Limitations, Artifacts, and Accuracy of Recovered Intercellular Stresses. PLoS One 8, e55172 (2013).

12. Nier, V. et al. Inference of Internal Stress in a Cell Monolayer. Biophys J 110, 1625– 1635 (2016).

13. Ng, M. R., Besser, A., Brugge, J. S. & Danuser, G. Mapping the dynamics of force transduction at cell-cell junctions of epithelial clusters. Elife 3, e03282 (2014).

14. Maruthamuthu, V., Sabass, B., Schwarz, U. S. & Gardel, M. L. Cell-ECM traction force modulates endogenous tension at cell-cell contacts. Proceedings of the National Academy of Sciences 108, 4708–4713 (2011).

15. McCain, M. L., Lee, H., Aratyn-Schaus, Y., Kléber, A. G. & Parker, K. K. Cooperative coupling of cell-matrix and cell-cell adhesions in cardiac muscle. Proc Natl Acad Sci U S A 109, 9881–9886 (2012).

16. Sim, J. Y. et al. Spatial distribution of cell-cell and cell-ECM adhesions regulates force balance while main taining E-cadherin molecular tension in cell pairs. Mol Biol Cell 26, 2456–2465 (2015).

17. Ruppel, A. et al. Force propagation between epithelial cells depends on active coupling and mechano-structural polarization. Elife 12, 1–42 (2023).

18. Trepat, X. et al. Physical forces during collective cell migration. Nat Phys 5, 426–430 (2009).

19. Balasubramaniam, L. et al. Investigating the nature of active forces in tissues reveals how contractile cells can form extensile monolayers. Nat Mater 20, 1156–1166 (2021).

20. Vincent, R. et al. Active Tensile Modulus of an Epithelial Monolayer. Phys Rev Lett 115, (2015).

21. Serra-Picamal, X. et al. Mechanical waves during tissue expansion. Nat Phys 8, 628– 634 (2012).

22. Haas, A. J. et al. Interplay between Extracellular Matrix Stiffness and JAM-A Regulates Mechanical Load on ZO-1 and Tight Junction Assembly. Cell Rep 32, 107924 (2020).

23. Mohan, A. et al. Spatial Proliferation of Epithelial Cells Is Regulated by E-Cadherin Force. Biophys J 115, 853–864 (2018).

24. Gayrard, C., Bernaudin, C., Déjardin, T., Seiler, C. & Borghi, N. Src- and confinement-dependent FAK activation causes E-cadherin relaxation and β-catenin activity. Journal of Cell Biology 217, 1063–1077 (2018).

25. Bonnet, I. et al. Mechanical state, material properties and continuous description of an epithelial tissue. J R Soc Interface 9, 2614–2623 (2012).

26. Bambardekar, K., Clément, R., Blanc, O., Chardès, C. & Lenne, P.-F. Direct laser manipulation reveals the mechanics of cell contacts in vivo. Proceedings of the National Academy of Sciences 112, 1416–1421 (2015).

27. Davis, J. R. et al. ECM degradation in the Drosophila abdominal epidermis initiates tissue growth that ceases with rapid cell-cycle exit. Current Biology 32, 1285–1300.e4 (2022).

28. Sonam, S. et al. Mechanical stress driven by rigidity sensing governs epithelial stability. Nat Phys 19, 132–141 (2023).

29. Vazquez, K., Saraswathibhatla, A. & Notbohm, J. Effect of substrate stiffness on friction in collective cell migration. Sci Rep 12, (2022).

30. Goodwin, K. et al. Basal Cell-Extracellular Matrix Adhesion Regulates Force Transmission during Tissue Morphogenesis. Dev Cell 39, 611–625 (2016).

31. Green, Y., Fredberg, J. J. & Butler, J. P. Relationship between velocities, tractions, and intercellular stresses in the migrating epithelial monolayer. Phys Rev E 101, (2020).

32. Friedrich, B. M. & Safran, S. A. How cells feel their substrate: Spontaneous symmetry breaking of active surface stresses. Soft Matter 8, 3223–3230 (2012).

33. Yang, W., Luo, M., Gao, Y., Feng, X. & Chen, J. Mechanosensing model of fibroblast cells adhered on a substrate with varying stiffness and thickness. J Mech Phys Solids 171, (2023).

34. Mertz, A. F. et al. Scaling of Traction Forces with the Size of Cohesive Cell Colonies. Phys Rev Lett 108, 198101 (2012).

35. Banerjee, S. & Marchetti, M. C. Contractile Stresses in Cohesive Cell Layers on Finite-Thickness Substrates. Phys Rev Lett 109, 108101 (2012).

36. Edwards, C. M. & Schwarz, U. S. Force localization in contracting cell layers. Phys Rev Lett 107, 1–5 (2011).

37. Prost, J., Jülicher, F. & Joanny, J. F. Active gel physics. Nat Phys 11, 111–117 (2015).

38. Oakes, P. W., Banerjee, S., Marchetti, M. C. & Gardel, M. L. Geometry regulates traction stresses in adherent cells. Biophys J 107, 825–833 (2014).

39. Solowiej-Wedderburn, J. & Dunlop, C. M. Sticking around: Cell adhesion patterning for energy minimization and substrate mechanosensing. Biophys J 121, 1777–1786 (2022).

40. Reinhart-King, C. A., Dembo, M. & Hammer, D. A. Cell-cell mechanical communication through compliant substrates. Biophys J 95, 6044–6051 (2008).

41. Kozyrina, A. N., Piskova, T. & Di Russo, J. Mechanobiology of Epithelia From the Perspective of Extracellular Matrix Heterogeneity. Front Bioeng Biotechnol 8, (2020).

42. Reichmann, E., Ball, R., Groner, B. & Friis, R. R. New mammary epithelial and fibroblastic cell clones in coculture form structures competent to differentiate functionally. J Cell Biol 108, 1127–1138 (1989).

43. Guvendiren, M. & Burdick, J. A. Stiffening hydrogels to probe short- and long-term cellular responses to dynamic mechanics. Nat Commun 3, 792 (2012).

44. Lopez, J. I., Kang, I., You, W. K., McDonald, D. M. & Weaver, V. M. In situ force mapping of mammary gland transformation. Integrative Biology 3, 910–921 (2011).

45. Maschler, S. M. et al. Tumor cell invasiveness correlates with changes in integrin expression and localization. Oncogene 24, 2032–2041 (2005).

46. Loebel, C., Mauck, R. L. & Burdick, J. A. Local nascent protein deposition and remodelling guide mesenchymal stromal cell mechanosensing and fate in three-dimensional hydrogels. Nat Mater 18, 883–891 (2019).

47. Loebel, C. et al. Metabolic labeling of secreted matrix to investigate cell–material interactions in tissue engineering and mechanobiology. Nature Protocols vol. 17 618– 648 Preprint at 10.1038/s41596-021-00652-9 (2022).

48. Kwon, M. Y. et al. Influence of hyaluronic acid modification on CD44 binding towards the design of hydrogel biomaterials. Biomaterials 222, 119451 (2019).

49. Kechagia, J. Z., Ivaska, J. & Roca-Cusachs, P. Integrins as biomechanical sensors of the microenvironment. Nat Rev Mol Cell Biol 20, 457–473 (2019).

50. Han, S. J., Bielawski, K. S., Ting, L. H., Rodriguez, M. L. & Sniadecki, N. J. Decoupling substrate stiffness, spread area, and micropost density: A close spatial relationship between traction forces and focal adhesions. Biophys J 103, 640–648 (2012).

51. Fusco, S., Panzetta, V. & Netti, P. A. Mechanosensing of substrate stiffness regulates focal adhesions dynamics in cell. Meccanica 52, 3389–3398 (2017).

52. Panzetta, V., Fusco, S. & Netti, P. A. Cell mechanosensing is regulated by substrate strain energy rather than stiffness. Proc Natl Acad Sci U S A 116, 22004–22013 (2019).

53. Han, S. J., Oak, Y., Groisman, A. & Danuser, G. Traction microscopy to identify force modulation in subresolution adhesions. Nat Methods 12, 653–656 (2015).

54. Solowiej-Wedderburn, J. & Dunlop, C. M. Cell-strain-energy costs of active control of contractility. Phys Rev E 107, (2023).

55. Mertz, A. F. et al. Cadherin-based intercellular adhesions organize epithelial cell-matrix traction forces. Proc Natl Acad Sci U S A 110, 842–847 (2013).

56. Rheinlaender, J. et al. Cortical cell stiffness is independent of substrate mechanics. Nat Mater 19, 1019–1025 (2020).

57. Valon, L., Marín-Llauradó, A., Wyatt, T., Charras, G. & Trepat, X. Optogenetic control of cellular forces and mechanotransduction. Nat Commun 8, (2017).

58. Doss, B. L. et al. Cell response to substrate rigidity is regulated by active and passive cytoskeletal stress. Proc Natl Acad Sci U S A 117, 12817–12825 (2020).

59. Ayad, N. M. E., Kaushik, S. & Weaver, V. M. Tissue mechanics, an important regulator of development and disease. Philosophical Transactions of the Royal Society B: Biological Sciences vol. 374 Preprint at 10.1098/rstb.2018.0215 (2019).

60. Bazellières, E. et al. Control of cell-cell forces and collective cell dynamics by the intercellular adhesome. Nat Cell Biol 17, 409–420 (2015).

61. Palmquist, K. H. et al. Reciprocal cell-ECM dynamics generate supracellular fluidity underlying spontaneous follicle patterning. Cell 185, 1960–1973.e11 (2022).

62. De Mets, R. et al. Cellular tension encodes local Src-dependent differential β1 and β3 integrin mobility. Mol Biol Cell 30, 181–190 (2019).

63. Von Erlach, T. C., Hedegaard, M. A. B. & Stevens, M. M. High resolution Raman spectroscopy mapping of stem cell micropatterns. Analyst 140, 1798–1803 (2015).

64. Pérez-González, C. et al. Active wetting of epithelial tissues. Nat Phys 15, 79–88 (2019).

65. Wang, Y. C. The origin and the mechanism of mechanical polarity during epithelial folding. Seminars in Cell and Developmental Biology vol. 120 94–107 Preprint at 10.1016/j.semcdb.2021.05.027 (2021).

66. Sharrock, T. E. & Sanson, B. Cell sorting and morphogenesis in early Drosophila embryos. Semin Cell Dev Biol 107, 147–160 (2020).

67. Chen, T., Saw, T. B., Mege, R. M. & Ladoux, B. Mechanical forces in cell monolayers. J Cell Sci 131, (2018).

68. Yamada, S. & Nelson, W. J. Localized zones of Rho and Rac activities drive initiation and expansion of epithelial cell-cell adhesion. Journal of Cell Biology 178, 517–527 (2007).

