## Supplemental information for "Emergence of active contractile patterns alters monolayer force generation and transmission in response to focal adhesion distribution"

#### Active stress model for confluent cell layers

The active stress model for the cell layer is based on a continuum description with active stress incorporated into the constitutive relations. This is a widely adopted approach, for a detailed treatment see e.g. [1, 2]. Here, due to the relative thinness of the cell layer compared with its extent, and the lack of cell rearrangements (see Extended Data Fig. 1(l)), we assume plane stress and a (linear) elastic response. We additionally assume that the within layer contractility has no preferred direction (isotropic contraction), to give the constitutive relations

$$\sigma_{ij} = \frac{hE}{1+\nu}\left(\epsilon_{ij} + \frac{\nu}{1-\nu}\epsilon_{kk}\delta_{ij}\right) + \frac{hE}{2(1-\nu)}P(\mathbf{x})\delta_{ij}, \tag{1}$$

where $E$ is the Young's modulus of the cells, $\nu$ the Poisson's ratio of the cells, and $h$ the thickness of the layer. The function $P(\mathbf{x})$ describes the contractility throughout the cell layer and we initially assume a uniform contractility throughout so that $P(\mathbf{x}) = P_0$. Later where we consider non-uniform contractility, we require a contractility function $P$ such that being closer to the edge of the layer means greater contractile activity. In this case, we follow [3, 4] and use the functional form $P(r) = a(1 + br^5)$. Although any monotonic increasing function would be sufficient [3], a polynomial form has the added benefit that in circular geometries (as considered here) explicit analytical solutions may be obtained. The degree $n = 5$ is chosen to ensure a sufficiently sharp transition from the edge region into centre where there is a more uniform contractile activity. The parameter $a$ is a scaling parameter for the total contractility of the layer. The parameter $b$ represents the extent of the localisation of that contractility to the edge of the layer. See Fig. 1 for some example radial profiles of differential contractility for different parameters $a$ and $b$.

The model is completed through a force balance equation. Here, we follow [1], and take

$$\nabla \cdot \sigma - KT(\mathbf{x})\mathbf{u} = \mathbf{0}, \tag{2}$$

where $T(\mathbf{x})$ is an indicator function that denotes discrete regions of adhesion so that $T(\mathbf{x}) = 1$ where the layer is adhered and $T(\mathbf{x}) = 0$ where there is no adhesion. The boundary conditions are zero stress on the outer boundary and continuity of both stress and deformation at all internal boundaries. The substrate resistance to deformation has been approximated as linear in the deformations (as if it were comprised of a dense network of linear springs). This approximation becomes exact for thin or stiff gels where deformations become localised, see e.g. the papers [2, 5]. The parameter $K$ thus quantifies the stiffness of the substrate and its resistance to deformation.

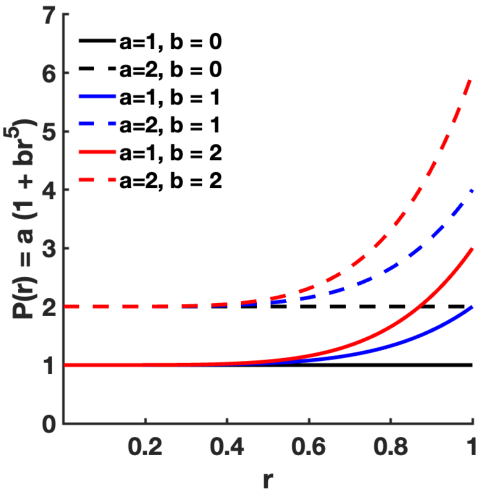


Figure 1: Sketch of differential contractility $P(r) = a(1 + br^5)$. Underlying contractility $a$ is taken as 1 (solid lines) or 2 (dashed lines) each illustrating the effect of varying localisation factors $b$.

#### Fixing model parameters

In the constitutive relation (1) we take the parameters as per Table 1. The key control parameter determining model solutions is the non-dimensional parameter $\gamma^2 = K(1-\nu^2)r_0^2/hE$, [1, 4], which quantifies the relative stiffness of substrate and cell layer. The other key element is the contractility function $P$. $K$ describes the stiffness of the substrate and for very thin substrates it can be shown analytically that this parameter is proportional to Young's modulus of the gel $E_s$, i.e. $K = \kappa E_s$ [2]. We make this approximation for the substrates used here, validating the approximation from the traction force data. Specifically, for complete adhesion we see that integrating the force balance over the domain $A$ and applying the boundary conditions gives $K = \int_A \nabla \cdot \sigma \mathrm{d}A / \int_A \mathbf{u} \mathrm{d}A$. In Fig. 2, we plot the data from the disc traction force measurements $K = \Sigma T/\Sigma u$ against the gel stiffness $E_s$. We observe an approximately linear relationship validating the approximation. Taking the mean ratio of $K/E_s$ from all the traction force data gave $\kappa$ as per Table 1. Finally where the contractility is constant so that $P = P_0$, $P_0$ may be fitted to either the strain energy or traction data collected from the traction force measurements, and we select traction throughout, see below.

#### Computational domains

The experiments are performed on two adhesion patterns: a completely functionalised disk (radius $100\mu$m) and a functionalised annular ring with inner and outer radii, $80\mu$m and $100\mu$m, respectively. For the disk we assume complete adhesion so that $T(\mathbf{x}) \equiv 1$ across the whole domain $|\mathbf{r}|/r_0 < 1$. For the ring of outer radius $100\mu$m and inner radius $80\mu$m we consider two patterns of adhesion. For the phase plane analysis, see the main paper Figures 4(a) and 4(b), we consider the ideal arrangement of $T(\mathbf{x}) \equiv 1$ in $0.8 < |\mathbf{r}|/r_0 < 1$, taking advantage of the analytical solutions this symmetry affords (see below). For all other simulations we consider a pattern of adhesion informed by the experimental pFAK imaging data (see e.g. main paper Fig. 1(h)). Specifically, we consider two thin annuli of thickness $r_t = 0.06$ with $T(\mathbf{x}) \equiv 1$ in $0.8 < |\mathbf{r}|/r_0 < 0.8 + r_t$ and $1 - r_t < |\mathbf{r}|/r_0 < 1$, corresponding to the regions with highest intensity pFAK. Additionally, we set a distribution of adhered spots in

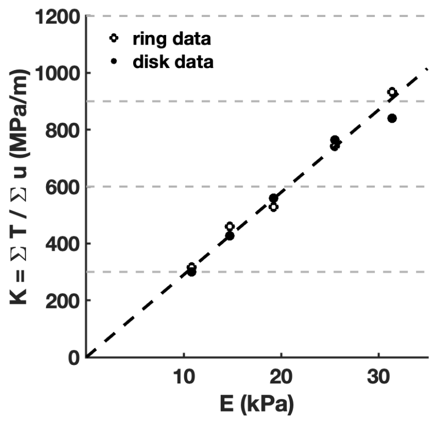


Figure 2: Traction to deformation ratio scales linearly with gel stiffness. Plot shows the ratio of the sum of tractions over the sum of displacements as derived from the experimental data plotted against Young's modulus ($E$). The dashed line has slope $\kappa$, as per Table 1, the mean ratio from all experimental data.

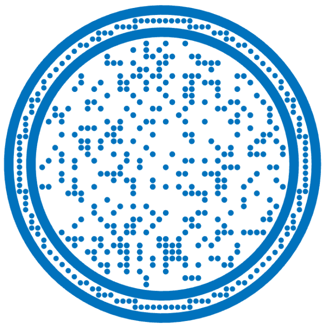

Figure 3: Computational adhesion pattern for simulations of adhesive rings.

| Parameter | Value | Source |
|---|---|---|
| Young's modulus: cell $E$ | 1kPa | literature see e.g. [6] |
| Layer thickness $h$ | 9$\mu$m | measured (Fig. 1 k and l in the main paper) |
| Poisson's ratio: cell $\nu$ | 0.45 | within literature range: 0.38–0.5 [7, 8] |
| Domain size: radius $r_0$ | 100$\mu$m | adhesive pattern size |
| Substrate parameter: $\kappa$ | 29000m$^{-1}$ | estimated from traction data |

Table 1: Constitutive parameter values used to calculate dimensionless parameter $\gamma^2 = K(1-\nu^2)r_0^2/hE$ which is used to quantify the relative stiffness of the substrate to the monolayer throughout the study.

the internal region and between the two annuli as the pFAK intensity is non-zero in these regions; spots between the two annuli have a radius of $r_s = 0.0167$ and in the internal region have radius $r_s = 0.018$. The pattern of spot placement used throughout is shown in Fig. 3.

**Sensitivity analysis of adhesion patterning**

To explore how sensitive the results are to the precise patterning of adhesion, and the possible role of imperfect adhesion in functionalised patches, we simulated different adhesive patterns and looked at their effect on the reported quantities of total traction and internal radial stress at ablation point. For the disk we compared the completely adhered disk used in the study to a simulation with discrete adhesions randomly distributed across the disk, as per Fig. 4A. Specifically, a square grid was generated across the disk such that there were 50 grid squares across the diameter. Adhesions with radius $0.0179r_0$ were then placed in randomly selected grid squares such that the total covered area is approximately 30% the total area of the disk. On this domain, solving on the random domain for the deformations with $\gamma = 15$ and uniform contractility, we calculate the total traction to be $\approx 93\%$ of the total traction calculated with complete adhesion across the entire interface. With the same parameters, we observed similarly small changes in the radial stress at ablation ($\sigma_{rr}(r = 0.5r_0)$): we calculate the $\sigma_{rr}(r = 0.5r_0)$ on the random domain to be $\approx 97\%$ of $\sigma_{rr}(r = 0.5r_0)$ of a disk with complete adhesion.

We further compare different adhesion patterns for the annular micropatterns. The annular micropattern is engineered to cover a region 80$\mu$m$< r <$ 100$\mu$m such that rings have a width of 20$\mu$m; this could be idealised to an adhesion arrangement with $T(\mathbf{x}) \equiv 1$ in $0.8 < |\mathbf{r}|/r_0 < 1$ (Fig. 4B). In table 2 we show that for such ideal arrangements a $\pm10\mu$m change in internal radius of the disk makes at most a 3% difference to the total traction measured across the layer and similarly small (4%) differences to the radial stress at the point of ablation. Indeed, even with a reduction in adhered area to $r_1 = 0.95$ we see a $\leq 10\%$ change in the total traction and radial stress. For smaller adhered areas we begin to see a sharper drop in the model predicted total traction and radial stress (see e.g. $r_1 = 0.975$ in table 2). As the arrangement of adhesion throughout the cell layer is non-uniform, we also consider different adhesion patterns inspired by the pFAK coverage (see main paper Fig. 1(j)). Specifically, we considered adhesion patterns with two thin annuli width $r_t$ such that $T(\mathbf{x}) \equiv 1$ in $0.8 < |\mathbf{r}|/r_0 < 0.8 + r_t$ and $1 - r_t < |\mathbf{r}|/r_0 < 1$ (as in Fig. 4C) to represent the regions of higher intensity pFAK. Reducing the width of the two annuli resulted in a decrease in both the tractions transmitted to the gel and internal radial stress, however, even when the adhesion percentage was reduced by 61% (from 36% of the disk with $r_1 = 0.8$ to 14% with $r_t = 0.04$), the reduction in total traction and internal stress was stil only 4%. We further considered a non-uniform pattern of adhered spots distributed between the two annuli with $r_t = 0.06$ (Fig. 4D), where spots had radius $r_s = 0.0167$; the spots were selected from the same square grid that was used in Fig. 4A to fill the desired domain. This addition of adhesions between the two annuli results in traction and stress values that lie between the cases of completely adhered annulus and two annuli without spots, as expected. Finally, we include a selection of spots randomly selected from a square grid with 40 squares across the diameter of the internal region as shown in Fig. 3. This addition of internal spots makes only 1% difference to the total traction and internal stress calculated from the model simulations, despite increasing the adhered area from 26% to 37% of the whole surface area of the cell layer.

| | Adhesion percentage | $T_{total}$ | $T_{total}$ of $T_{total}(r_1 = 0.8)$ | $\sigma_{rr}(r = 0.5)$ | $\sigma_{rr}(r = 0.5)$ of $\sigma_{rr}(r = 0.5; r_1 = 0.8)$ |
|---|---|---|---|---|---|
| RINGS, e.g. Fig. 4B | | | | | |
| $r_1 = 0.7$ | 51% | 4.39 | 101% | 0.72 | 101% |
| $r_1 = 0.75$ | 44% | 4.38 | 100% | 0.72 | 101% |
| $r_1 = 0.8$ | 36% | 4.37 | 100% | 0.72 | 100% |
| $r_1 = 0.85$ | 28% | 4.33 | 99% | 0.71 | 99% |
| $r_1 = 0.9$ | 19% | 4.25 | 97% | 0.69 | 96% |
| $r_1 = 0.95$ | 10% | 4.01 | 92% | 0.65 | 90% |
| $r_1 = 0.975$ | 5% | 3.60 | 83% | 0.58 | 81% |
| TWO THIN RINGS, e.g. Fig. 4C | | | | | |
| $r_t = 0.04$ | 14% | 4.18 | 96% | 0.69 | 96% |
| $r_t = 0.06$ | 22% | 4.29 | 98% | 0.70 | 98% |
| $r_t = 0.08$ | 29% | 4.34 | 99% | 0.71 | 99% |
| TWO THIN RINGS WITH SPOTS | | | | | |
| Fig. 4D | 26% | 4.31 | 99% | 0.71 | 99% |
| Fig. 3 | 37% | 4.35 | 100% | 0.72 | 100% |
| Fig. 4E | 62% | 4.36 | 100% | 0.72 | 101% |

Table 2: Sensitivity analysis of different adhesion simulations of annular adhesion patterned gel with contractility maintained at $P_0 = 1$. Total traction, $T_{total}$, and radial stress at point of ablation, $\sigma_{rr}(r = 0.5)$, are computed for different adhesion patterns (shown in Fig. 4) on a gel with stiffness $\gamma = 15$. The values are also given relative to an annulus with internal radius $r_1 = 0.8$ simulating perfect adhesion across the micropattern. Adhesion percentage denotes the proportion of the circular geometry that is 'adhered'.

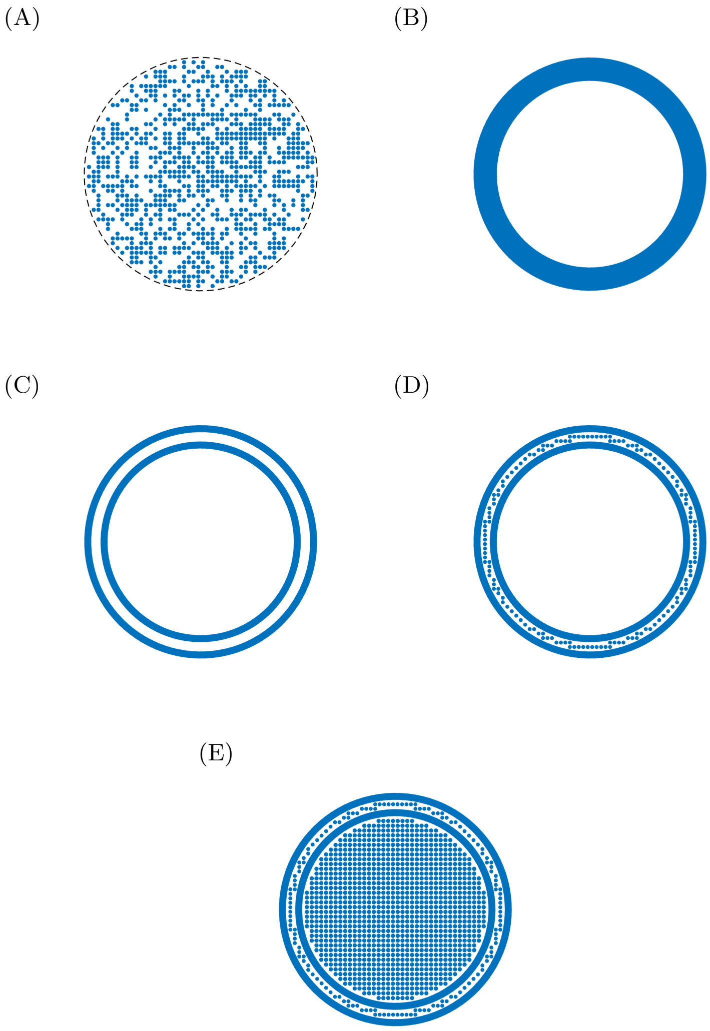


Figure 4: Computational adhesive patterns used for sensitivity analysis: (A) shows a distribution of 935 adhesive spots randomly selected from a uniform grid across a circular geometry for an adhesion coverage of 30%; (B) shows adhesion across an annular region. The internal radius of the annulus is $r_1 \times r_0$, where $r_0$ is the external radius; here $r_1 = 0.8$ is depicted; (C) shows adhesion across two annuli for a circular geometry. Both annuli have width $r_t \times r_0$, here the inner-most radius is $r_1 = 0.8$ and $r_t = 0.06$; (D) shows adhesion across the same geometry as (C) with additional adhesive spots between the two annuli. These spots have radius $r_s = 0.017$; (E) shows adhesion across the same geometry as (D) with additional adhesive spots in a square grid with 40 grid squares across the diameter of the internal region. These spots have radius $r_s = 0.018$.

#### Solving for deformations

##### Analytical solutions for complete adhesion on disk and ring

For an adhered disk, $T(\mathbf{x}) \equiv 1$ across $|\mathbf{r}| < r_0$ analytical solutions of (2) can be obtained with both $P = P_0$ and $P(r) = a(1 + br^5)$, see [4]. In both cases, symmetry implies the deformations will be purely radial so that $\mathbf{u} = u(r)\mathbf{e_r}$. For $P = P_0$,

$$\frac{u}{r_0} = \frac{-P_0(1+\nu)}{2\left(\gamma I_0(\gamma) - (1-\nu)I_1(\gamma)\right)} I_1\left(\frac{\gamma r}{r_0}\right), \tag{3}$$

where $I_1$ is the modified Bessel function and $\gamma^2 = K(1-\nu^2)r_0^2/hE$. For the radially varying profile we obtain

$$\frac{u(r/r_0)}{r_0} = \frac{(1+\nu)}{2\gamma}\left(cI_1\left(\frac{\gamma r}{r_0}\right) + \frac{5ab}{\gamma^5}\left[\gamma^4 r^4 + 15\gamma^2 r^2 - \frac{45\pi}{2}L_1\left(\frac{\gamma r}{r_0}\right)\right]\right), \tag{4}$$

where we have defined

$$c := \frac{-a(1+b)}{F(\gamma)} + \frac{5ab}{\gamma^5 F(\gamma)}\left[(1-\nu)\left(\gamma^3 + 15\gamma - \frac{45\pi}{2\gamma}L_1(\gamma)\right) - \left(5\gamma^3 + 45\gamma - \frac{45\pi}{2}L_0(\gamma)\right)\right],$$

given in terms of the modified Struve functions $L_0(z)$ and $L_1(z)$ [9]. Note we use non-dimensional scaled spatial variables here (i.e. introducing $r/r_0$) to enable solutions to be domain size independent, in this formulation $b$ is now also scaled to be non-dimensional so that $b = b^*/r_0^5$ (where $b^* \equiv b$ in dimensional variables). Function $F(\gamma)$ is defined by

$$F(z) := I_0(z) + \frac{(\nu - 1)}{z}I_1(z).$$

Similarly analytical solutions for both contractility profiles ($P = P_0$ and $P(r) = a(1 + br^5)$) can be obtained for the adhered ring pattern, provided symmetry is maintained and no distributed adhesions spots are assumed. However, in this case the solutions are more mathematically complicated involving additional special functions. See [4] for the full details of these solutions and their derivations. We note that the parameter $\gamma^2 = K(1-\nu^2)r_0^2/hE$ continues to be the control parameter in the analytical solutions.

##### Numerical solutions for pFAK informed adhesion patterns

When considering adhesions distributed in discrete regions, such as the adhesion geometry shown in Fig. 3, numerical solutions must be obtained using finite element methods. We again work on the non-dimensional domain scaled to have maximum radius 1, as such we continue to interpret parameter $b$ in the non-dimensional sense as for the analytic solutions. For each defined geometry, a mesh was generated automatically using the generateMesh command within the MATLAB PDE Toolbox (with `Hmax`, maximum edge length, set at 0.005). The force balance equation may be expressed as a general elliptic PDE in two dimensions

$$\begin{aligned} -\nabla \cdot (c_{11}\nabla u_1) - \nabla \cdot (c_{12}\nabla u_2) + a_{11}u_1 + a_{12}u_2 &= f_1 \\ -\nabla \cdot (c_{21}\nabla u_1) - \nabla \cdot (c_{22}\nabla u_2) + a_{21}u_1 + a_{22}u_2 &= f_2. \end{aligned}$$

As in the radially symmetric cases of a complete adhesion or an adhered ring, we have normalised length scales by the cell radius $r_0$. Thus, specifically the coefficient matrices are

$$c_{11} = \begin{pmatrix} 1 & 0 \\ 0 & \frac{(1-\nu)}{2} \end{pmatrix}, \quad c_{12} = \begin{pmatrix} 0 & \nu \\ \frac{(1-\nu)}{2} & 0 \end{pmatrix}, \quad c_{21} = \begin{pmatrix} 0 & \frac{(1-\nu)}{2} \\ \nu & 0 \end{pmatrix}, \quad c_{22} = \begin{pmatrix} \frac{(1-\nu)}{2} & 0 \\ 0 & 1 \end{pmatrix},$$

$(a_{11}, a_{22}) = T(\mathbf{x})\gamma^2$, and $a_{12} = a_{21} = 0$. With differential contractility given by $P(r) = a(1 + br^5)$, the coefficients on the right-hand-side are given by $f_1 = 5((1+\nu)/2)abx\left(x^2 + y^2\right)^{3/2}$ and $f_2 = 5((1+\nu)/2)aby\left(x^2 + y^2\right)^{3/2}$, where we have used $r = \left(x^2 + y^2\right)^{1/2}$ to convert between Cartesian and polar coordinates. The no stress boundary condition is input as generalised Neumann boundary conditions

$$\begin{aligned} \mathbf{n} \cdot (c_{11} \nabla u_1) + \mathbf{n} \cdot (c_{12} \nabla u_2) + q_{11} u_1 + q_{12} u_2 &= g_1 \\ \mathbf{n} \cdot (c_{21} \nabla u_1) + \mathbf{n} \cdot (c_{22} \nabla u_2) + q_{21} u_1 + q_{22} u_2 &= g_2, \end{aligned}$$

with $\mathbf{n}$ the normal vector to the boundary; $q_{11} = q_{12} = q_{21} = q_{22} = 0$ and $(g_1, g_2) = -((1 + \nu)/2)a\left(1 + b(x^2 + y^2)^{5/2}\right)\mathbf{n}$. To obtain solutions for a uniform contractility throughout the epithelium, we set $a = P_0$ and $b = 0$.

### Parameter fitting to experimental total traction data

From the previous section, we can see that for a given adhesion geometry the model is completely parameterised by $\gamma$ and $\nu$ with either $P_0$ defined (uniform contractility) or $a$ and $b$ defined (radially varying contractility). The parameters $\gamma$ and $\nu$ are determined from the mechanical properties of the gel and epithelium as described above. The Poisson's ratio for all cells is expected to be close to 0.5 (the value for an incompressible material) and little variation is expected, the specific value 0.45 is taken within the range commonly reported for cells (see table 1).

We used the measured total traction data from the traction force microscopy to set the contractility parameter $P_0$ or $a$ (for a given choice of $b$) for each gel stiffness. Specifically for each contractility model (uniform or varying) the model deformations $\mathbf{u}$ are derived as described above and from these deformations the total applied tractions $T_{tot}$ are calculated

$$T_{tot}(P(\mathbf{x})) = \int_A KT(\mathbf{x})\mathbf{u} dA. \tag{5}$$

$T_{tot}$ is then fit to the experimental data of $\Sigma T$ via $T_{tot} = \Sigma T \cdot A_{pix}$, where $A_{pix}$ is the area of a pixel (here $0.519\mu$m$\times 0.519\mu$m).

As an alternative to fitting to the traction data we could have chosen to fit to the total strain energy data. However, as the strain energy data is a derived quantity from the traction force data we chose to fit our single fit parameter to the traction directly. The model with these contractility parameters then subsequently predicts qualitatively similar trends in the strain energy as observed (see e.g. Figs. 4(g,l)), however, on rings, at a slightly higher level than observed. One contributory factor to the observed difference in strain energy is the approximation of a linear spring resistance in the force balance (2), which on finite depth gels and at low adhesion percentages may overpredict strain energy due to the contribution of non-local effects.

When fitting the model with differential contractility, the localisation of contractility to the edge of the layer, $b$, also needs to be assigned. We choose $b$ to fit the trends in recoil velocity so that the relative drop in internal stress matches the relative drop in recoil velocity. We assume recoil velocity and internal radial stress are linearly correlated but cannot fit $b$ directly to the measured velocities. In considering relative changes we arbitrarily select the starting values for the localisation factor $b$. We set values of $b = 2.1$ and $b = 0.1$ on the softest gels for disk and ring adhesion patterns, respectively. From these initial parameters choices, we then set $b$ values on each stiffer gel so that the relative drop of internal stress matches the same drop in the recoil velocity.

To illustrate this process in more detail consider the contours of constant total traction and constant radial stress shown in Fig. 5 for $\gamma = 5$. For our first softest substrate a range of pairs of $a$ and $b$ could be chosen to fit any particular total traction data, for example the two points highlighted (black crosses) have the same total traction. However, once the arbitrary reference value radial stress for this substrate is selected then the choice is constrained (as e.g. although the two crosses have the same total traction they represent different radial stress values). For each subsequent stiffness substrate simulated there is only only one point in parameter space that can satisfy both the requirements of fitting the observed traction data and the required internal stress value (this will be the single intersection of the two relevant contours in the parameter space). For example, in Fig. 5 although we have indicated two parameter choices with black crosses which correspond to the same total traction, the values of internal stress for these two parameter choices are different. Our initial reference value is only constrained by the requirements that the total contractility and concentration of contractility are not unfeasible. We additionally must choose initial values that leave enough of the parameter space open to fit the data on stiffer substrates.

In the main paper we discuss the need for spatial distributions of contractility to emerge in order to explain the observed traction stresses and recoil velocities. We illustrated this using parameter phase planes Figures 4a and b (main paper) chosen for particular values of $b$, and contours of total contractility and internal radial stress. We finish this section by further plotting additional contours across a range of parameter values, Fig. 6, to demonstrate that the contour lines of constant total traction and internal radial stress depicted in the main paper show general trends predicted by the model. We see that all pairs of contours plotted in this sweep demonstrate the same qualitative features as discussed in the paper and that the selected values were representative.

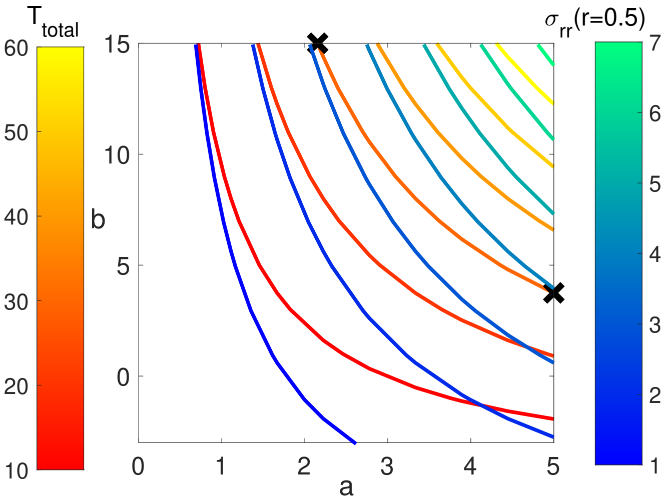


Figure 5: Contour plot shows lines of constant total traction (red-orange) and constant internal stress (blue-green) as $a$ and $b$ are varied. Black crosses mark two choices of $a$ and $b$ parameters such that $T_{total}$ is the same at both points, however, the value of internal stress ($\sigma_{rr}(r = 0.5)$ is different at each point). Here substrate stiffness is kept fixed at $\gamma = 5$.

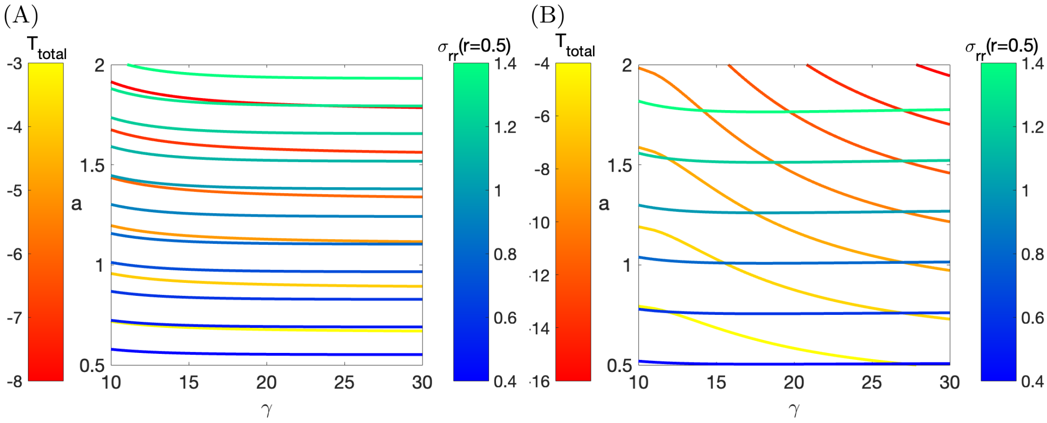


Figure 6: Contour plot shows lines of constant total traction (red-orange) and constant internal radial stress (blue-green) as $a$ is varied on different stiffness substrates. (A) Contractility is uniform ($b = 0$): here the traction and stress contours follow similar trajectories. (B) Contractility is localised to the edge of the layer ($b = 1$): here there is a much greater overlap in traction and stress contour lines.

### References

[1] Josephine Solowiej-Wedderburn and Carina. M. Dunlop. Sticking around: Cell adhesion patterning for energy minimization and substrate mechanosensing. *Biophys J*, 121(9):1777–1786, 2022.

[2] Shiladitya Banerjee and M. Cristina Marchetti. Contractile stresses in cohesive cell layers on finite-thickness substrates. *Phys. Rev. Lett.*, 109(10):108101, 2012.

[3] Carina Dunlop. Differential cellular contractility as a mechanism for stiffness sensing. *New Journal of Physics*, 21(6):063005, 2019.

[4] Josephine Solowiej-Wedderburn and Carina M Dunlop. Cell-strain-energy costs of active control of contractility. *Physical review E*, 107(6):L062401, 2023.

[5] F. Mertz, Aaron, S. Benerjee, Yonglu Che, Guy K. German, Ye Xu, Callen Hyland, M. Cristina Marchetti, Valerie Horsley, and Eric. R. Dufresne. Scaling of traction forces with the size of cohesive colonies. *Phys. Rev. Lett.*, 108(19):198101, 2012.

[6] Jordi Alcaraz, Ren Xu, Hidetoshi Mori, Celeste M Nelson, Rana Mroue, Virginia A Spencer, Doug Brownfield, Derek C Radisky, Carlos Bustamante, and Mina J Bissell. Laminin and biomimetic extracellular elasticity enhance functional differentiation in mammary epithelia. *The EMBO journal*, 27(21):2829–2838, 2008.

[7] J Rheinlaender, A Dimitracopoulos, B Wallmeyer, N M Kronenberg, K J Chalut, M C Gather, T Betz, G Charras, and K Franze. Cortical cell stiffness is independent of substrate mechanics. *Nat Mater*, 19(9):1019–1025, 2020.

[8] CA Mullen, TJ Vaughan, MC Voisin, MA Brennan, P Layrolle, and LM McNamara. Cell morphology and focal adhesion location alters internal cell stress. *J R Soc Interface*, 11(101):20140885, 2014.

[9] M Abramowitz and I A Stegun. *Handbook of mathematical functions: with formulas, graphs, and mathematical tables*, volume 55. Courier Corporation, 1964.

### Supplementary Table 1: Main figures statistical values

| Figure | N numbers (gels) | | | | | Unit | Mean | | | | | Standard Deviation | | | | | Kruskal-Wallis | Mann-Whitney | | | |
|---|---|---|---|---|---|---|---|---|---|---|---|---|---|---|---|---|---|---|---|---|---|
| | 10 | 15 | 20 | 25 | 30 | | 10 | 15 | 20 | 25 | 30 | 10 | 15 | 20 | 25 | 30 | | 15 | 20 | 25 | 30 |
| 1i | 20 (5) | 23 (6) | 30 (8) | 25 (5) | 20 (4) | N | 122.5 | 109.6 | 150.9 | 104.4 | 116.7 | 23.1 | 23.3 | 50.9 | 34.0 | 26.8 | 0.006 | 0.113 | 0.122 | 0.098 | 0.394 |
| 1j | 28 (11) | 29 (11) | 23 (7) | 23 (6) | 28 (7) | N | 126.5 | 109.2 | 122.1 | 129.5 | 137.2 | 31.2 | 20.9 | 34.5 | 33.2 | 42.6 | 0.048 | 0.049 | 0.454 | 0.747 | 0.476 |
| 2c | 20 (5) | 23 (6) | 30 (8) | 25 (5) | 20 (4) | μm | 4.08 | 2.47 | 1.18 | 1.01 | 0.67 | 1.95 | 2.02 | 0.87 | 0.51 | 0.58 | $1.4 \times 10^{-18}$ | $8.1 \times 10^{-4}$ | $1.3 \times 10^{-7}$ | $3.5 \times 10^{-8}$ | $2.2 \times 10^{-7}$ |
| 2d | 28 (11) | 29 (11) | 23 (7) | 23 (6) | 28 (7) | μm | 3.34 | 2.20 | 2.14 | 1.11 | 1.15 | 2.58 | 0.97 | 2.54 | 0.94 | 0.80 | $1.4 \times 10^{-10}$ | 0.123 | 0.003 | 1.9 x 10-6 | $6.5 \times 10^{-7}$ |
| 2e | 20 (5) | 23 (6) | 30 (8) | 25 (5) | 20 (4) | Pa | $8.44 \times 10^{7}$ | $4.82 \times 10^{7}$ | $3.51 \times 10^{7}$ | $4.56 \times 10^{7}$ | $2.69 \times 10^{7}$ | $4.35 \times 10^{7}$ | $1.56 \times 10^{7}$ | $2.00 \times 10^{7}$ | $2.22 \times 10^{7}$ | $1.07 \times 10^{7}$ | $4.8 \times 10^{-9}$ | 0.002 | $2.2 \times 10^{-5}$ | 0.001 | $2.0 \times 10^{-6}$ |
| 2f | 28 (11) | 29 (11) | 23 (7) | 23 (6) | 28 (7) | Pa | $5.75 \times 10^{7}$ | $5.13 \times 10^{7}$ | $5.23 \times 10^{7}$ | $4.78 \times 10^{7}$ | $5.91 \times 10^{7}$ | $5.23 \times 10^{7}$ | $2.25 \times 10^{7}$ | $3.15 \times 10^{7}$ | $3.15 \times 10^{7}$ | $2.77 \times 10^{7}$ | 0.255 | 0.434 | 0.684 | 0.421 | 0.070 |
| 2g | 20 (5) | 23 (6) | 30 (8) | 25 (5) | 20 (4) | Pa | 1859.0 | 1655.1 | 894.1 | 1170.6 | 706.5 | 824.9 | 1171.9 | 645.3 | 853.0 | 541.7 | $3.0 \times 10^{-8}$ | 0.116 | $3.9 \times 10^{-5}$ | 0.001 | $2.7 \times 10^{-6}$ |
| 2h | 28 (11) | 29 (11) | 23 (7) | 23 (6) | 28 (7) | Pa | 1765.8 | 1630.1 | 1927.6 | 1210.7 | 1724.3 | 1491.2 | 1037.3 | 2326.7 | 1100.3 | 1694.7 | 0.137 | 0.842 | 0.201 | 0.044 | 0.787 |
| 2i | 20 (5) | 23 (6) | 30 (8) | 25 (5) | 20 (4) | pJ | 26399.7 | 8261.6 | 3126.2 | 3272.1 | 1157.6 | 24238.5 | 7927.5 | 3417.4 | 3089.4 | 1355.2 | $3.2 \times 10^{-14}$ | 0.004 | $1.6 \times 10^{-6}$ | $3.3 \times 10^{-6}$ | $3.0 \times 10^{-7}$ |
| 2j | 28 (11) | 29 (11) | 23 (7) | 23 (6) | 28 (7) | pJ | 19440.6 | 7893.4 | 9135.1 | 4291.8 | 5959.2 | 42344.8 | 6744.3 | 13299.0 | 5777.0 | 7889.4 | 0.001 | 0.660 | 0.057 | $9.6 \times 10^{-4}$ | 0.005 |
| 3k | 20 (5) | 23 (6) | 30 (8) | 25 (5) | 20 (4) | μm/s | 0.138 | 0.136 | 0.110 | 0.112 | 0.110 | 0.079 | 0.062 | 0.050 | 0.021 | 0.024 | 0.172 | 0.827 | 0.163 | 0.758 | 0.570 |
| 3l | 28 (11) | 29 (11) | 23 (7) | 23 (6) | 28 (7) | μm/s | 0.189 | 0.145 | 0.131 | 0.110 | 0.117 | 0.083 | 0.048 | 0.048 | 0.051 | 0.036 | $6.8 \times 10^{-6}$ | 0.037 | 0.004 | $5.5 \times 10^{-5}$ | $2.2 \times 10^{-4}$ |
| 3o | 20 (5) | 23 (6) | 30 (8) | 25 (5) | 20 (4) | Pa | $-1.79 \times 10^{7}$ | $-1.18 \times 10^{7}$ | $-0.67 \times 10^{7}$ | $-0.75 \times 10^{7}$ | $-0.39 \times 10^{7}$ | $1.46 \times 10^{7}$ | $0.81 \times 10^{7}$ | $0.83 \times 10^{7}$ | $0.42 \times 10^{7}$ | $0.27 \times 10^{7}$ | $1.8 \times 10^{-5}$ | 0.192 | 0.006 | 0.023 | 0.003 |
| 3p | 28 (11) | 29 (11) | 23 (7) | 23 (6) | 28 (7) | Pa | $-2.29 \times 10^{7}$ | $-1.27 \times 10^{7}$ | $-1.66 \times 10^{7}$ | $-0.98 \times 10^{7}$ | $-1.30 \times 10^{7}$ | $2.84 \times 10^{7}$ | $0.87 \times 10^{7}$ | $1.52 \times 10^{7}$ | $0.97 \times 10^{7}$ | $1.10 \times 10^{7}$ | 0.068 | 0.105 | 0.158 | 0.007 | 0.100 |
| 3q | 20 (5) | 23 (6) | 30 (8) | 25 (5) | 20 (4) | pJ | 10716.3 | 3348.6 | 1435.4 | 973.7 | -323.8 | 11383.4 | 5373.9 | 2772.3 | 726.9 | 2680.0 | $3.4 \times 10^{-10}$ | 0.021 | $2.3 \times 10^{-4}$ | $5.9 \times 10^{-4}$ | $1.0 \times 10^{-4}$ |
| 3r | 28 (11) | 29 (11) | 23 (7) | 23 (6) | 28 (7) | pJ | 14753.8 | 3660.9 | 3296.1 | 1548.7 | 2969.1 | 29915.0 | 4381.3 | 6933.5 | 2206.8 | 5415.6 | $3.4 \times 10^{-5}$ | 0.045 | 0.008 | $4.5 \times 10^{-5}$ | $9.3 \times 10^{-5}$ |
| 5b | 32 (3) | 22 (3) | 69 (3) | 48 (3) | 61 (3) | 16-bit Grey levels | 15173.1 | 15866.3 | 23751.7 | 10.974.1 | 12973.7 | 7123.3 | 9272.0 | 10696.3 | 5461.2 | 7753.0 | $7.9 \times 10^{-15}$ | 0.867 | $7.1 \times 10^{5}$ | 0.005 | 0.059 |
| 5c | 34 (4) | 55 (4) | 59 (3) | 62 (2) | 41 (2) | 16-bit Grey levels | 15836.4 | 15285.4 | 16551.1 | 15368.4 | 14684.5 | 8430.0 | 8312.1 | 6529.4 | 7101.4 | 10850 | 0.037 | 0.602 | 0.980 | 0.400 | 0.104 |
| 5d | 32 (3) | 22 (3) | 69 (3) | 48 (3) | 61 (3) | 16-bit Grey levels | 11168.0 | 11032.1 | 16890.4 | 11593.0 | 8938.0 | 5933.1 | 5240.4 | 5279.6 | 5762.0 | 4645.5 | $3.4 \times 10^{-19}$ | 0.979 | $2.1 \times 10^{-7}$ | 0.520 | 0.008 |

| | | | | | | | | | | | | | | | | | | | | |
|---|---|---|---|---|---|---|---|---|---|---|---|---|---|---|---|---|---|---|---|---|
| 5e | 24 (3) | 32 (3) | 59 (3) | 62 (2) | 41 (2) | 16-bit Grey levels | 16196.4 | 12414.3 | 8869.2 | 12372.9 | 9840.0 | 10533.9 | 6074.4 | 2161.9 | 3565.3 | 4245.2 | 5.0 $\times 10^{-12}$ | 0.048 | 1.0 $\times 10^{-6}$ | 0.081 | 1.9 $\times 10^{-4}$ |
| 5f | 32 (3) | 22 (3) | 69 (3) | 48 (3) | 61 (3) | 16-bit Grey levels | 15750.0 | 16459.8 | 15743.4 | 10402.9 | 11641.1 | 3847.4 | 5954.3 | 3990.1 | 3870.0 | 2873.5 | 5.4 $\times 10^{-17}$ | 0.965 | 0.899 | 5.1 $\times 10^{-7}$ | 1.1 $\times 10^{-6}$ |
| 5g | 34 (4) | 55 (4) | 59 (3) | 62 (2) | 41 (2) | 16-bit Grey levels | 28160.5 | 35148.9 | 18770.7 | 14541.5 | 17514.6 | 16042.0 | 12749.0 | 6013.8 | 3282.0 | 5099.7 | 7.2 $\times 10^{-26}$ | 0.011 | 0.020 | 3.1 $\times 10^{-6}$ | 0.002 |
| 5r (disk) | 48 (4) | 78 (3) | 95 (3) | 51 (2) | 58 (3) | % | 5.580 | 2.799 | 2.999 | 1.519 | 2.972 | 6.200 | 5.190 | <0.001 | 3.533 | 5.029 | 0.006 | 0.007 | 0.010 | 4.0 $\times 10^{-4}$ | 0.022 |
| 5r (ring) | 97 (2) | 21 (3) | 49 (3) | 28 (2) | 14 (3) | % | 1.441 | 1.648 | 1.201 | 2.391 | 6.720 | 1.657 | 2.721 | 2.324 | 3.363 | 3.511 | 2.6 X $10^{-6}$ | 0.482 | 0.466 | 0.154 | 2.9 $\times 10^{-7}$ |

Supplementary Table 2: Supplementary figures statistical values

| Figure | N numbers (gels) | | | | | | Mean | | | | | Standard Deviation | | | | | Kruskal-Wallis | Mann-Whitney | | | |
|---|---|---|---|---|---|---|---|---|---|---|---|---|---|---|---|---|---|---|---|---|---|
| | 10 | 15 | 20 | 25 | 30 | Unit | 10 | 15 | 20 | 25 | 30 | 10 | 15 | 20 | 25 | 30 | | 15 | 20 | 25 | 30 |
| S1c | 28 (7) | 28 (7) | 28 (7) | 28 (7) | 24 (6) | Pa | 10810.4 | 14667.5 | 19201.8 | 25494.1 | 31418.5 | 2563.2 | 1574.4 | 2696.9 | 3795.2 | 2794.8 | $1.2 \times 10^{-60}$ | $1.9 \times 10^{-7}$ | $2.7 \times 10^{-10}$ | $1.3 \times 10^{-10}$ | $6.5 \times 10^{-10}$ |
| S1d | 87 (3) | 56 (3) | 33 (3) | 40 (1) | 17 (2) | % | 24.0 | 23.2 | 21.2 | 19.6 | 20.0 | 3.5 | 2.3 | 2.1 | 2.1 | 3.0 | $3.3 \times 10^{-23}$ | 0.002 | $3.0 \times 10^{-8}$ | $3.4 \times 10^{-13}$ | $1.7 \times 10^{-6}$ |
| S1e | 41 (3) | 13 (3) | 36 (3) | 15 (2) | 41 (3) | % | 17.6 | 17.2 | 14.7 | 15.4 | 15.5 | 1.7 | 1.7 | 2.3 | 1.6 | 2.5 | $1.2 \times 10^{-9}$ | 0.342 | $1.3 \times 10^{-8}$ | $1.5 \times 10^{-4}$ | $7.8 \times 10^{-5}$ |
| S1j | 20 (5) | 23 (6) | 30 (8) | 25 (5) | 20 (4) | % | 0.32 | 0.22 | 0.11 | -0.13 | 0.53 | 0.90 | 0.71 | 1.01 | 0.71 | 1.29 | 0.227 | 0.970 | 0.945 | 0.123 | 0.560 |
| S1k | 28 (11) | 29 (11) | 23 (7) | 23 (6) | 28 (7) | % | 0.47 | 0.16 | -0.42 | 1.12 | 0.12 | 1.62 | 0.60 | 1.22 | 6.72 | 0.71 | 0.288 | 0.615 | 0.074 | 0.389 | 0.652 |
| S2g | 20 (5) | 23 (6) | 30 (8) | 25 (5) | 20 (4) | pixels | 542025 | 218209 | 121461 | 114767 | 61743 | 307498 | 84989 | 67864 | 46357 | 24890 | $1.3 \times 10^{-21}$ | $2.0 \times 10^{4}$ | $4.7 \times 10^{-7}$ | $5.3 \times 10^{-7}$ | $1.2 \times 10^{-7}$ |
| S2h | 28 (11) | 29 (11) | 23 (7) | 23 (6) | 28 (7) | pixels | 350437 | 215146 | 191101 | 124234 | 122272 | 324791 | 87074 | 125988 | 84106 | 55581 | $1.2 \times 10^{-11}$ | 0.031 | 0.003 | $2.8 \times 10^{-}$ | $6.7 \times 10^{-8}$ |
| S3i | 20 (5) | 23 (6) | 30 (8) | 25 (5) | 20 (4) | seconds | 26.2 | 16.9 | 20.6 | 18.6 | 22.2 | 295.3 | 81.2 | 45.7 | 36.5 | 111.7 | 0.705 | 0.336 | 0.317 | 0.174 | 0.457 |
| S3j | 28 (11) | 29 (11) | 23 (7) | 23 (6) | 28 (7) | seconds | 24.2 | 18.9 | 21.6 | 20.1 | 19.2 | 40.7 | 157.5 | 410.8 | 744.7 | 49.4 | 0.961 | 0.955 | 0.656 | 0.932 | 0.550 |
| S4l (disk) | 48 (4) | 78 (3) | 95 (3) | 51 (2) | 58 (3) | % | 7.802 | 1.069 | 3.046 | 2.015 | 2.484 | 9.625 | 8.382 | 8.927 | 6.205 | 7.925 | $4.3 \times 10^{-4}$ | $1.9 \times 10^{-4}$ | 0.002 | 0.002 | 0.002 |
| S4l (ring) | 97 (2) | 21 (3) | 49 (3) | 28 (2) | 14 (3) | % | 1.651 | 1.464 | 2.605 | 6.389 | 7.022 | 5.659 | 6.994 | 7.441 | 9.563 | 6.089 | 0.007 | 0.800 | 0.753 | 0.012 | 0.002 |
